# TDP2 couples PKA–CREB signalling to genome stability in human cerebellar development

**DOI:** 10.64898/2026.11.16.767189

**Authors:** Wanpeng Wang, Guoming Ma, Fei Wang, Yushan Deng, Yuqiu Zheng, Zhichao Qiu, Minmin Lou, Wanda Zhan, Jin Tao, Karen T. Tao, Huan Li, Jingjing Xie, Zhuobin Liang, Liemin Zhou, Yidi Sun, Wenyuan Wang

**Author notes:** These authors contributed equally: Wanpeng Wang, Guoming Ma, Fei Wang, Yushan Deng.

## Abstract

Transcription and neuronal activity in the central nervous system generate abundant DNA damage, particularly DNA double-strand breaks (DSBs), rendering neural cells — and cerebellar neurons especially — vulnerable to genome instability^1–5^. Tyrosyl-DNA phosphodiesterase 2 (TDP2, also known as TTRAP) resolves topoisomerase-2-linked DSBs, and biallelic TDP2 mutations cause spinocerebellar ataxia autosomal recessive 23 (SCAR23)^6,7^, yet how DNA repair shapes human cerebellar development is unknown. Here we model SCAR23 in cerebellar organoids derived from patient induced pluripotent stem cells carrying TDP2 c.650del and from prime-edited isogenic TDP2 c.400C>T lines. TDP2 inactivation caused organoid hypoplasia during FGF8b-dependent cerebellar specification, which was reversed by TDP2 complementation. Single-nucleus transcriptomics across three developmental stages revealed early dysregulation of ribosome biogenesis, attenuated CREB1 regulon activity and a subsequent failure of Purkinje cell maturation. TDP2 inactivation upregulated PKIB, a cerebellum-enriched inhibitor of protein kinase A, lowering PKA and CREB phosphorylation. Whole-genome sequencing showed excess de novo deletions in mutant organoids, enriched at candidate cis-regulatory elements of neurodevelopmental genes. Pharmacological activation of the cAMP–PKA cascade restored CREB phosphorylation, reduced DNA damage and rescued organoid growth. These findings link developmental signalling to genome maintenance in the human cerebellum and identify the cAMP–PKA–CREB axis as a candidate therapeutic target in DNA-repair-deficient cerebellar disease.

## Introduction

Genome instability, and the accumulation of DNA damage and mutations that follows from it, has been implicated in neuronal activity, neurodevelopment, neuronal migration and neurodegeneration, but its precise contribution to each remains unclear^1,8–11^. TDP2 repairs topoisomerase-2-induced DSBs and was independently identified as TTRAP and EAP2 through its interactions with members of the tumour necrosis factor receptor (TNFR) family, TNFR-associated factors (TRAFs) and ETS proteins^12–15^. Loss-of-function mutations in TDP2 cause spinocerebellar ataxia autosomal recessive 23 (SCAR23), a rare syndrome of severe cerebellar atrophy, ataxia, intellectual disability and drug-resistant epilepsy^16–22^, and TDP2 has also been implicated in sporadic and familial Parkinson’s disease^23,24^. Type II topoisomerases (TOP2) relieve topological stress but can be trapped during transcription and neuronal activity, leaving abortive TOP2–DSB complexes with 4-base-pair staggered ends^25–31^. TDP2 hydrolyses the phosphotyrosyl bond of these trapped adducts, a step thought to permit error-free religation^32^. Work to date has relied largely on Tdp2-knockout mice and on primary fibroblasts or lymphoblastoid lines from patients, neither of which recapitulates the SCAR23 phenotype; patient fibroblasts accumulate little DNA damage^33,34^. How loss of a ubiquitously expressed repair enzyme produces a cerebellum-restricted developmental failure, and what genomic error signatures it leaves during early human neurodevelopment, therefore remain open questions.

Here we established TDP2-mutant induced pluripotent stem cell (iPSC) lines from a Chinese SCAR23 family^21^ and differentiated them into human cerebellar organoids (hCerOs) to model the disease. Combining cellular assays, bulk RNA sequencing, single-nucleus transcriptomics and whole-genome sequencing (WGS), we examined the processes disrupted by TDP2 inactivation in iPSCs, iPSC-derived neural progenitor cells (iNPCs) and cerebellar organoids. Organoids derived from two TDP2-mutant lines acquired markedly more small insertions and deletions during differentiation than controls of nearly identical genetic background. We further find that TDP2 inactivation upregulates PKIB and attenuates PKA–CREB signalling, and that this hypoactivity accompanies cerebellar hypoplasia and genome instability during cerebellar specification. Together, these results identify a convergent molecular mechanism in SCAR23 and indicate a role for TDP2 in repair fidelity and genome maintenance during human cerebellar development.

## Results

### TDP2 inactivation drives cerebellar hypoplasia

SCAR23 presents with severe cerebellar atrophy on neuroimaging, whereas Tdp2-knockout mice reproduce neither the atrophy nor the ataxia. To define the developmental basis of this atrophy, we generated iPSCs from peripheral blood mononuclear cells (PBMCs) of a familial cohort^21^ comprising a patient homozygous for TDP2 c.650del with severe cerebellar atrophy and two healthy sisters, one wild-type and one heterozygous carrier (Fig. 1a and Extended Data Fig. 1a-c). We additionally engineered the pathogenic TDP2 c.400C>T mutation into wild-type iPSCs from the same family by CRISPR prime editing^35^ (Fig. 1a and Extended Data Fig. 1c). All lines showed normal morphology, expressed the pluripotency markers SOX2, LIN28A and SSEA4, and retained a normal karyotype (Extended Data Fig. 1b-f). Enzymatic assays confirmed complete loss of TDP2 catalytic activity in both homozygous mutants, whereas heterozygous c.650del cells retained partial activity (Fig. 1b and Extended Data Fig. 1g). TDP2 transcript levels were also reduced in both mutant lines (Fig. 1c).

**Fig. 1.**
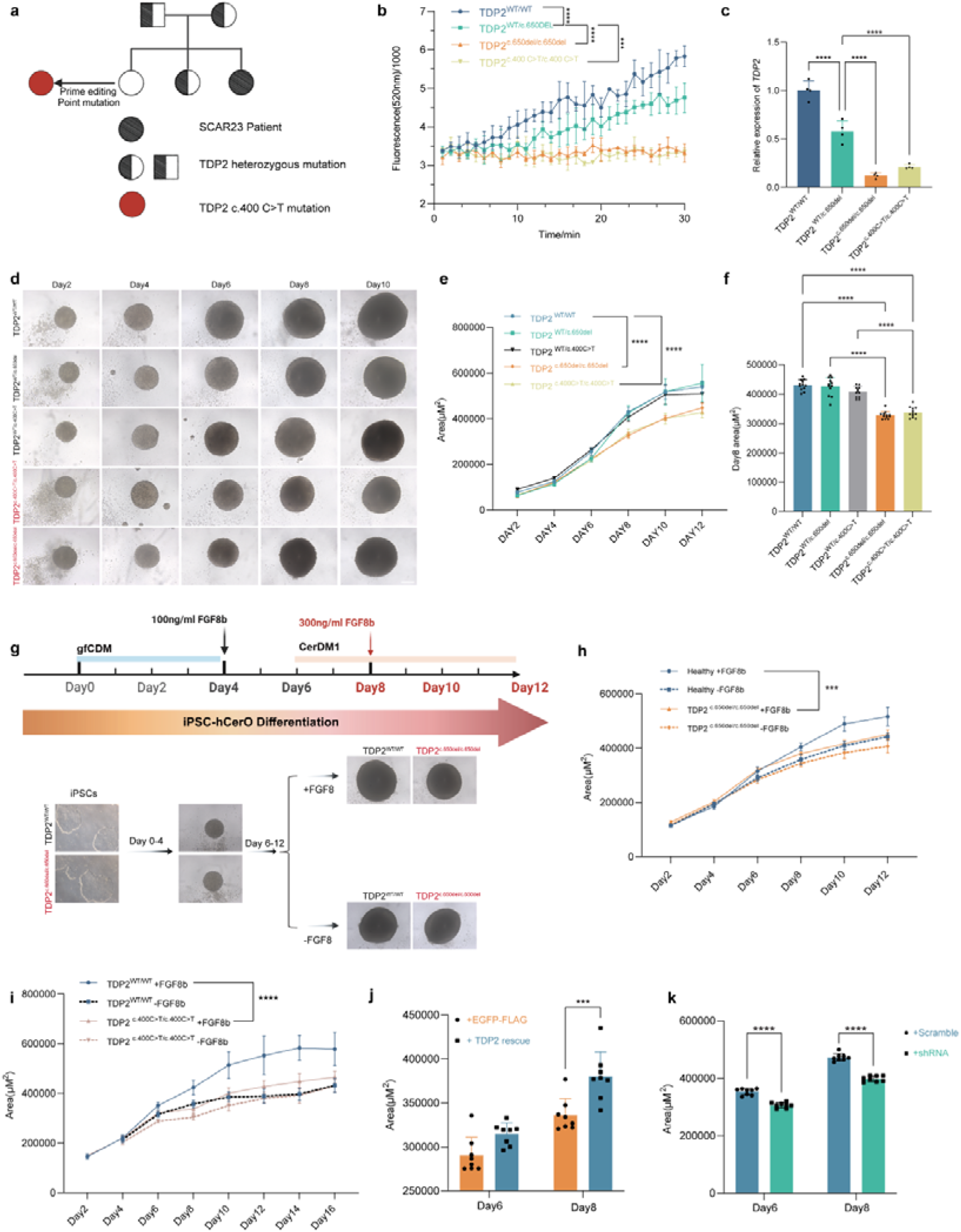
TDP2 inactivation impairs cerebellar specification and causes cerebellar organoid hypoplasia. **a,** Pedigree of the SCAR23 family with the indicated TDP2 mutation, and the prime-edited isogenic mutant line (TDP2 c.400C>T) generated from the healthy family member. **b,** Fluorescence-based biochemical assay of phosphodiesterase activity for wild-type and mutant TDP2 (mean ± s.d., n = 6 independent biological replicates). **c,** TDP2 transcript levels in wild-type and TDP2-mutant iPSC lines measured by RT–qPCR (mean ± s.d., n = 4). **d,** Representative bright-field images of human cerebellar organoids (hCerOs) derived from wild-type and TDP2-mutant lines. Scale bars, 200 μm. **e,f,** Cross-sectional area of hCerOs from day 0 to day 12 **(e)** and at day 8 **(f)** of induction (mean ± s.d., n = 12). **g,** Schematic of the hCerO differentiation protocol, indicating the stage of FGF8b administration required for cerebellar lineage commitment. **h,i,** hCerO area under standard and FGF8b-deprived conditions for the TDP2 c.650del **(h)** and TDP2 c.400C>T **(i)** lines compared with wild-type lines (mean ± s.d., n = 12). **j,** Exogenous expression of wild-type TDP2 in TDP2 c.650del iPSCs before hCerO induction; area of day 6 and day 8 hCerOs for control (EGFP–FLAG) and rescue (EGFP–TDP2) lines (mean ± s.d., n = 8). **k,** shRNA-mediated knockdown of TDP2 in wild-type iPSCs followed by hCerO induction; hCerO area for TDP2 shRNA and scrambled control lines (mean ± s.d., n = 8). Statistics: one-way ANOVA (b,c,f) or two-way ANOVA (e,h,i,j,k). ****P < 0.0001, ***P < 0.001, **P < 0.01, *P < 0.05.

We differentiated mutant and wild-type iPSCs into human cerebellar organoids (hCerOs) using an established protocol^36,37^. Mutant hCerOs showed pronounced volumetric hypoplasia from day 8 of differentiation, which persisted through mid and late stages, relative to three healthy control lines (one sibling from the SCAR23 family and two unrelated donors) and two heterozygous lines (Fig. 1d-f and Extended Data Fig. 2a-d). The onset of hypoplasia coincided with day 8, the stage at which elevated FGF8b signalling drives cerebellar specification, and withdrawal of FGF8b abolished the difference in organoid size between mutant and control lines (Fig. 1g-i and Extended Data Fig. 2e-g). TDP2 activity is therefore required during the FGF8b-dependent stage of cerebellar specification, and its loss produces a lineage-restricted arrest of early cerebellar patterning.

To test whether these defects were caused by loss of TDP2 itself, we generated shRNA knockdown lines from wild-type iPSCs and rescue lines in which wild-type TDP2 was expressed from a lentiviral vector in the c.650del background (Extended Data Fig. 3a). Knockdown in wild-type cells reproduced the early-onset hypoplasia, whereas complementation of the mutant line restored hCerO growth (Fig. 1j,k and Extended Data Fig. 3b-e). Pharmacological inhibition of TDP2 with TND1288 in combination with etoposide likewise recapitulated the hypoplastic phenotype (Extended Data Fig. 3f,g). TDP2 inactivation is thus intrinsically sufficient to impair cerebellar organogenesis, mirroring the clinical presentation of SCAR23.

### TDP2 inactivation dysregulates cAMP–PKA–CREB signalling from the earliest stages

To resolve how TDP2 inactivation reshapes human cerebellar development in a stage- and cell-type-specific manner, we performed single-nucleus RNA sequencing (snRNA-seq) of hCerOs at day 12 (D12), day 36 (D36) and day 63 (D63) (Fig. 2a). The dataset comprised wild-type (WT), heterozygous carrier (Het) and patient-derived homozygous TDP2 c.650del (Homo) organoids, together with the independently generated TDP2 c.400C>T (c400) line at D12 (Fig. 2a). After quality control and sample integration, high-quality nuclei were retained for downstream analyses (Extended Data Fig. 4a). By integrating our data with published single-cell RNA-seq data from human cerebellar organoids^36^ and snRNA-seq data from human neural organoids^38^ (Extended Data Fig. 4b,c), we assigned cells to 10 major types on the basis of known markers and regional identity: early progenitors (Pro), granule cell progenitors (GCP), interneuron progenitors (IP), PAX2+ interneuron progenitors (PIP), Purkinje-lineage populations (PC), Bergmann glia (BG), oligodendrocyte progenitor cells (OPC), endothelial cells (Endo), roof-plate cells (RP) and meninges (Extended Data Fig. 4d,e). Their temporal distribution captured the expected shift from progenitor-dominated states at D12 to increasingly diverse Purkinje-lineage and glial populations at D36 and D63 (Extended Data Fig. 4f). CytoTRACE2 assigned the highest predicted developmental potency to progenitors, consistent with their early identity, whereas most PC states occupied more differentiated positions (Extended Data Fig. 4g). Subclustering resolved 24 transcriptionally distinct subtypes (Fig. 2b and Extended Data Fig. 4h).

**Fig. 2.**
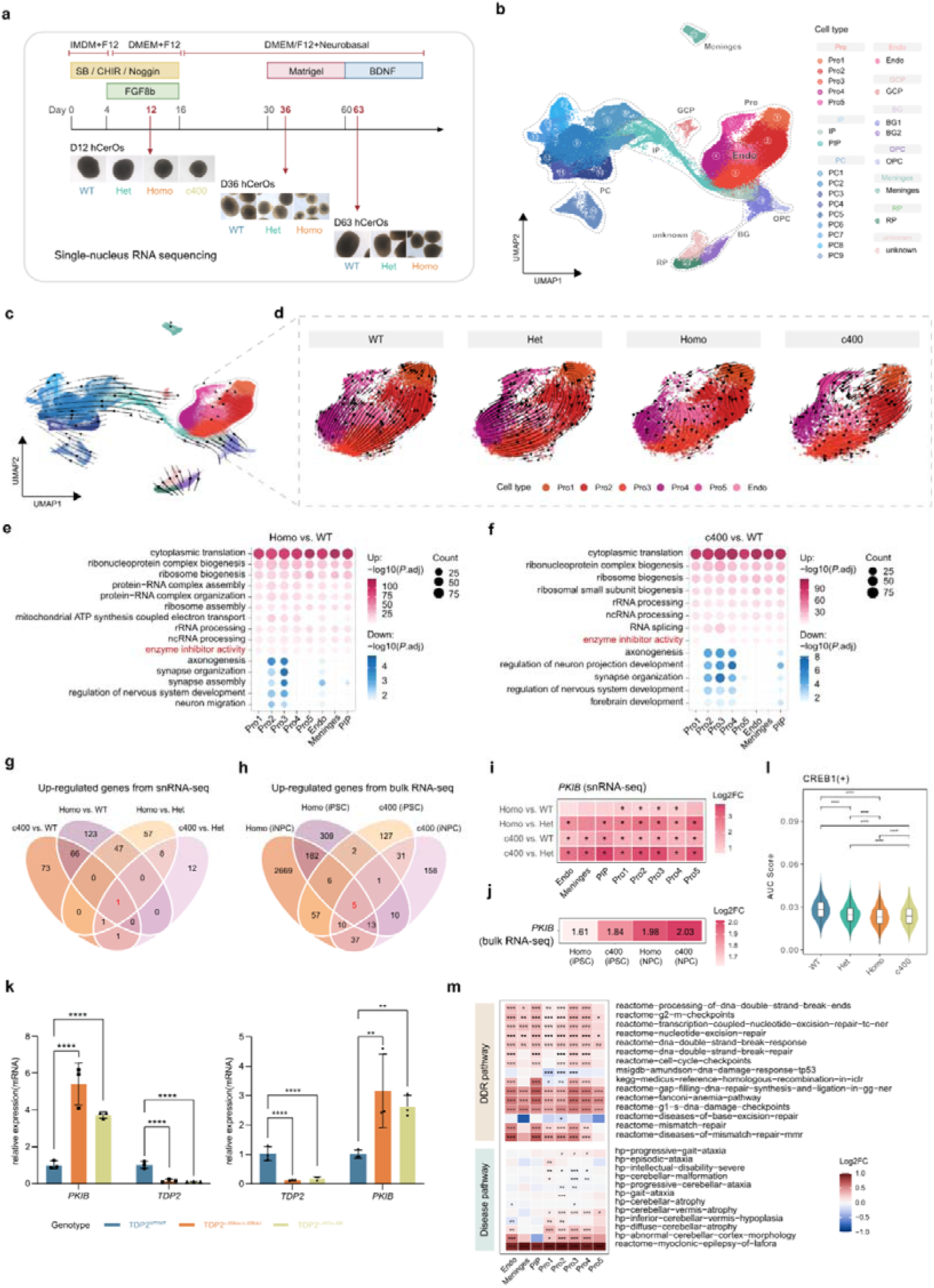
TDP2 inactivation dysregulates cAMP–PKA–CREB signalling and DNA repair gene programmes at D12. **a,** Schematic representation of hCerO differentiation and single-nucleus RNA sequencing (snRNA-seq) sampling timeline. WT, wild type; Het, heterozygous TDP2 c.650del; Homo, homozygous patient-derived TDP2 c.650del mutant; c400, prime-edited homozygous TDP2 c.400C>T line. **b,** Uniform manifold approximation and projection (UMAP) plot of integrated snRNA-seq data of hCerOs at day 12 (D12), day 36 (D36) and day 63 (D63) coloured by 24 cellular subtypes. **c,** RNA-velocity streamline of all cells. **d,** Stream plot of the RNA-velocity showing the difference across WT, Het, Homo, and c400 genotypes at D12. **e,f,** Dot plots showing enriched Gene Ontology (GO) terms for differentially expressed genes (DEGs) in Homo versus WT **(e)** and in c400 versus WT **(f)** at D12. **g,** Venn diagrams showing the overlap of upregulated DEGs (|log2 fold change| > 1 in at least two cell types at D12) across four comparisons (Homo versus WT, c400 versus WT, Homo versus Het, and c400 versus Het). **h,** Venn diagrams showing the overlap of upregulated DEGs (log2 fold change > 1) in TDP2-mutant lines (Homo and c400) relative to WT controls across iPSCs and iNPCs by bulk RNA sequencing (RNA-seq). **i,** Heatmap showing differential expression of PKIB in D12 cell populations. Asterisks indicate statistically significant comparisons. **j,** Heatmap shows log2 fold change of PKIB expression in TDP2-mutant iPSCs and iNPCs compared to WT controls. **k,** RT-qPCR quantification of TDP2 and PKIB expression in iNPCs and hCerOs (mean ± SD, n = 3). **l,** Violin plots showing CREB1(+) regulon activity in progenitor populations across genotypes. **m,** Heatmap of log2 fold change in curated DNA-damage-response and DNA-repair pathway scores (top) and cerebellar-disease associated pathway scores (bottom) between Homo and WT cells at D12. Red and blue denote higher and lower scores in Homo, respectively. Asterisks indicate Benjamini-Hochberg-adjusted Wilcoxon P-values (*P.adj < 0.05, **P.adj < 0.01, and ***P.adj < 0.001).

Given the early developmental abnormalities associated with TDP2 inactivation, we next asked whether they were accompanied by changes in the differentiation commitment of cerebellar cell populations. RNA-velocity streamlines broadly followed the expected trajectory from progenitors towards Purkinje-lineage populations, although directionality was less clearly defined within the early progenitor compartment (Fig. 2c). At D12, Homo and c400 progenitors showed markedly altered velocity vector orientations compared with WT and Het progenitors, including locally opposing vectors and a less coherent flow field (Fig. 2d). Because these alterations were shared by two independently derived TDP2-mutant lines, they indicate that predicted transcriptional state transitions are perturbed in TDP2-mutant progenitors before distinct PC states emerge.

To uncover the molecular programmes underlying these altered transcriptional dynamics, we identified differentially expressed genes (DEGs) between genotypes at D12. In Homo hCerOs, upregulated DEGs were enriched for cytoplasmic translation, ribosome biogenesis, ribosome assembly and related biosynthetic programmes across Pro1-Pro5, PIP, endothelial cells and meninges (Fig. 2e). Upregulated DEGs in c400 hCerOs were enriched for the same ribosome-associated and RNA-processing pathways (Fig. 2f), in line with previous evidence implicating TDP2 in ribosomal RNA biogenesis^23^. By contrast, downregulated DEGs in both mutant lines were associated with axonogenesis, synapse organization, regulation of nervous system development and neuronal migration (Fig. 2e,f). GO enrichment of Homo-versus-Het DEGs followed the same direction as the Homo-versus-WT comparison (Extended Data Fig. 4i). Because the two mutant lines share these changes but not their genetic background, the coordinated disruption of RNA biosynthesis and early neuronal programmes is attributable to TDP2 inactivation rather than to the family-specific background of the patient-derived line.

To trace the developmental onset of this dysregulation, and to ask whether *PKIB* upregulation preceded organoid differentiation, we examined bulk RNA sequencing (RNA-seq) data from earlier stages — undifferentiated iPSCs and iPSC-derived neural progenitor cells (iNPCs) (Extended Data Fig. 5a,b). Intersecting the DEGs of c.650del and c.400C>T iPSCs and iNPCs identified five genes upregulated more than twofold at every stage in both mutants: *PKIB, GTF2IP23, GPR85, TPH1* and *AMH* (Fig. 2h,j). Within D12 organoids, one gene *PKIB* was upregulated more than twofold across more than two cell types in both the Homo and c400 lines relative to WT and Het controls (Fig. 2g,i). *PKIB* was the only gene common to both analyses. *PKIB* encodes a cerebellum-enriched inhibitor of the cAMP–PKA pathway that suppresses phosphorylation of PKA and of its downstream targets, including CREB, a transcription factor required for cerebellar neuronal differentiation and survival^39–42^, and it was upregulated in iPSCs, iNPCs and D12 hCerOs of both mutant lines as well as in TDP2 knockdown lines (Fig. 2j,k and Extended Data Fig. 5c). PKIB upregulation is thus an early and reproducible correlate of TDP2 inactivation. Consistent with this, SCENIC analysis showed lower CREB1(+) regulon activity in Homo and c400 progenitors than in WT and Het progenitors (Fig. 2l and Extended Data Fig. 4j), indicating that CREB-dependent transcription is already attenuated in mutant progenitors.

We further asked whether these early alterations extended to gene programmes associated with TDP2-mediated DNA repair and with SCAR23-related cerebellar phenotypes. Cell-type-resolved rank-based gene set enrichment analysis at D12 revealed elevated scores for DNA damage response and repair (DDR) gene sets, including DSB response, transcription-coupled nucleotide excision repair, Fanconi anaemia, cell cycle checkpoint and mismatch repair (MMR) signatures, together with gene sets associated with diffuse cerebellar atrophy, cerebellar malformation and progressive gait ataxia (Fig. 2m).

Together these findings implicate dysregulation of the PKIB–PKA–CREB axis as an early molecular event following TDP2 inactivation, potentially linking altered transcriptional programmes to the DNA damage response and to impaired cerebellar organoid development. Read alongside the enrichment results, early PKIB upregulation did not mark accelerated differentiation; it coincided instead with sustained progenitor-associated RNA biosynthesis and loss of neuronal programmes, consistent with a stress-adapted but developmentally constrained state.

### TDP2-dependent cAMP–PKA–CREB signalling and genome stability in cerebellar development

TDP2 is required to repair DSBs caused by TOP2 trapping, but how it engages these lesions in neural cells is not well defined. To follow the TDP2 response to DSBs, we induced breaks with the TOP2-trapping agent etoposide (ETO). In SH-SY5Y neuroblastoma cells, ETO caused a modest increase in TDP2 abundance and rapid recruitment of TDP2 to damage sites, where it formed foci (Extended Data Fig. 6a-d). Live-cell imaging confirmed a damage-dependent increase in TDP2 foci (Extended Data Fig. 6e and Supplementary Videos 1 and 2). Doxorubicin, a second anthracycline that traps TOP2^43^, also induced TDP2 condensation at lesion sites (Extended Data Fig. 6f,g). By contrast, TDP2 was not recruited to TOP1-trapped lesions generated by camptothecin, which are processed by TDP1 (Extended Data Fig. 6h,i). TDP2 is therefore recruited selectively to TOP2-associated DSBs, where it forms foci.

TOP2-mediated DSBs carry ‘junk ends’, in which the break end is covalently linked to a tyrosine residue and requires processing by the proteasome, MRE11, Artemis and TDP2 before ligation (Extended Data Fig. 6j). To ask whether TDP2 is also recruited to clean-ended DSBs, we used a tetracycline-inducible AsiSI endonuclease system that generates DSBs at genomic sites separated by more than 1 Mb^44^. TDP2 was robustly recruited and formed foci at AsiSI-induced clean-ended lesions in SH-SY5Y cells (Extended Data Fig. 6k,l).

We next examined the consequences of TDP2 inactivation in the two patient-derived mutant iPSC lines, in which TDP2 expression was significantly reduced (Fig. 1c). Although these lines were no more sensitive to etoposide than wild-type controls, their DSB repair was markedly less efficient, as judged by γH2AX accumulation (Fig. 3a,b and Extended Data Fig. 7a). Mutant iPSCs also retained substantial numbers of micronuclei 24 h after damage^33,34,45,46^, a phenotype abolished by exogenous expression of wild-type TDP2 (Fig. 3c,d and Extended Data Fig. 7b). TDP2 is therefore required for efficient repair of TOP2-associated DSBs, and its loss compromises genome stability in iPSCs.

**Fig. 3.**
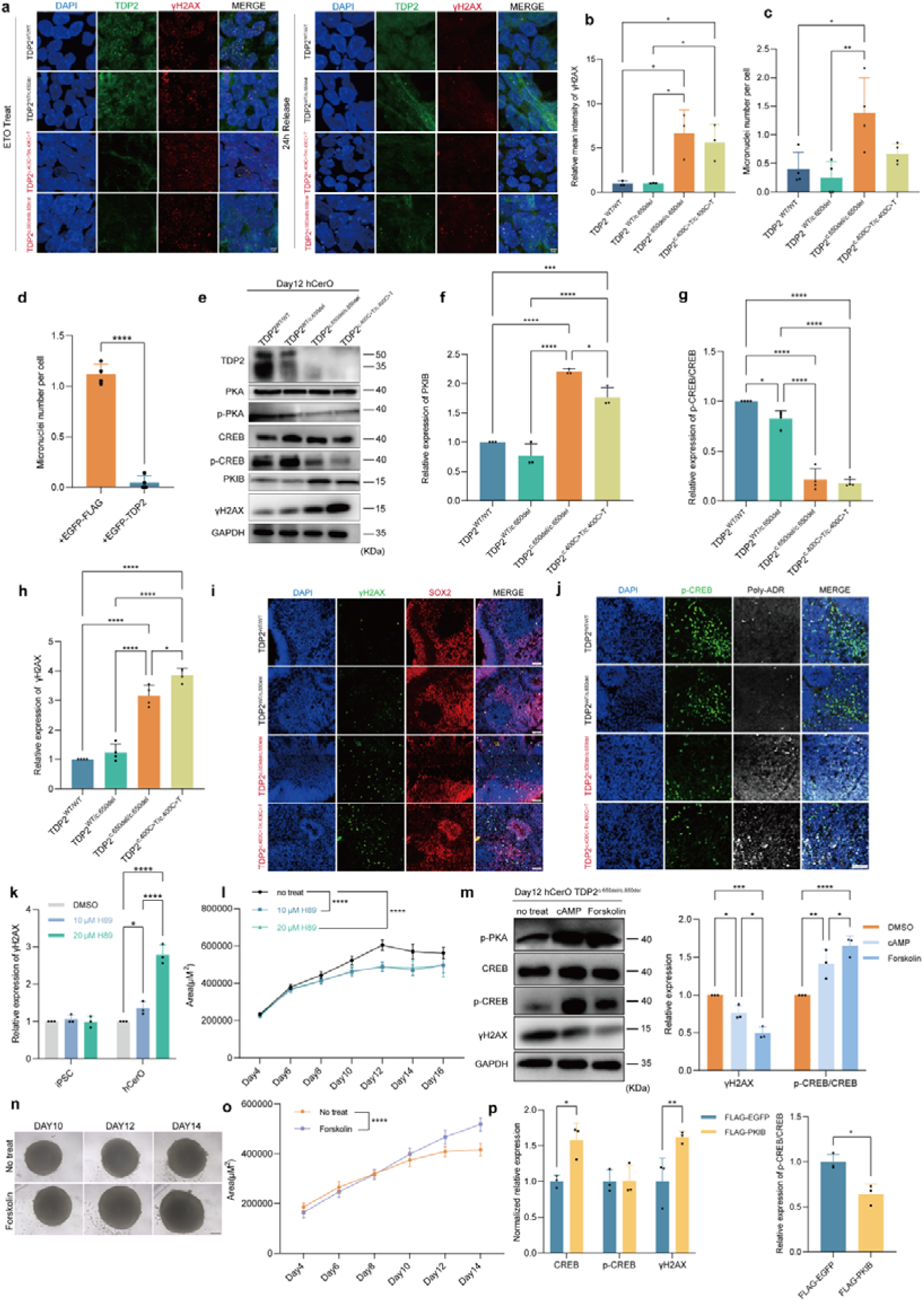
TDP2 acts through the cAMP–PKA–CREB cascade to preserve genome stability and developmental integrity during cerebellar ontogeny. **a,b,** Representative immunofluorescence **(a)** and quantification **(b)** of γH2AX intensity in iPSCs after etoposide (ETO) treatment and 24 h of recovery (mean ± s.d., n = 4). **c,** Micronucleus counts after ETO treatment and 24 h of recovery in iPSCs (mean ± s.d., n = 4). **d,** Micronucleus counts in TDP2 c.650del iPSCs overexpressing FLAG or FLAG–TDP2 after ETO treatment and 24 h of recovery (mean ± s.d., n = 4). Scale bars, 25 μm. **e,** Representative western blots of day 12 hCerOs for PKA–CREB pathway proteins and γH2AX. **f-h,** Band intensities for PKIB **(f)**, p-CREB/CREB **(g)** and γH2AX **(h)** in day 12 hCerOs, normalized to control (mean ± s.d., n = 4 for p-CREB and γH2AX; n = 3 for PKIB). **i,j,** Representative immunofluorescence of γH2AX with SOX2 **(i)** and of p-CREB with poly ADP-ribose **(j)** in day 12 wild-type and TDP2-mutant hCerOs. Scale bars, 50 μm. **k,** γH2AX band intensity in day 12 hCerOs treated with H89, normalized to control (mean ± s.d., n = 3). **l,** Cross-sectional area of wild-type hCerOs with and without H89 treatment (mean ± s.d., n = 12). **m,** Representative western blots of day 12 TDP2 c.650del hCerOs treated with the PKA agonists cAMP and forskolin, with band intensities for p-CREB/CREB and γH2AX (mean ± s.d., n = 3). **n,o,** Representative bright-field images (**n**; scale bars, 200 μm) and cross-sectional area **(o)** of TDP2 c.650del hCerOs with and without forskolin (mean ± s.d., n = 12). **p,** Band intensities for CREB, p-CREB, γH2AX and the p-CREB/CREB ratio in day 12 hCerOs overexpressing PKIB (mean ± s.d., n = 3). Statistics: one-way ANOVA (b,c,d,f,g,h) or two-way ANOVA (k,l,m,o,p). ****P < 0.0001, ***P < 0.001, **P < 0.01, *P < 0.05.

Abnormal protein aggregation is a feature of many neurodegenerative diseases, including Parkinson’s disease, Alzheimer’s disease and amyotrophic lateral sclerosis (ALS)^47^. We next asked how SCAR23-causative mutations affect TDP2 focus dynamics. Exogenous expression of the c.650del or c.454G>A mutants, but not c.400C>T, produced distinct pathological aggregates in HEK293T cells (Extended Data Fig. 7c,d), and aberrant aggregation of endogenous c.650del protein was also seen in pre-permeabilized iPSCs (Extended Data Fig. 7e). Structural modelling with AlphaFold 3 and analysis of intrinsically disordered regions (IDRs) indicated that these mutations do not introduce additional IDRs^48^ (Extended Data Fig. 7f,g). C-terminal EGFP tagging of wild-type TDP2 did not affect nuclear localization but abolished focus assembly (Extended Data Fig. 7h,i). Because the C-terminal domain mediates catalysis and DNA binding (Extended Data Fig. 1d), these results suggest that impaired DNA binding in this region contributes to the aberrant condensation of the c.650del protein.

Human cells express two topoisomerase II isoforms, TOP2α and TOP2β, which relieve topological stress during replication and transcription, respectively^49^. To ask whether TDP2 responds to DSBs arising from both processes, we synchronized SH-SY5Y cells in G1 and treated them with ETO^50,51^ (Extended Data Fig. 8a). TDP2 formed prominent foci that colocalized with γH2AX specifically in EdU-negative G1 cells, whereas condensation was nearly absent in EdU-positive S/G2 cells (Extended Data Fig. 8b). The same G1-restricted condensation was observed in iNPCs (Extended Data Fig. 8c,d). Inhibiting transcription with 5,6-dichlorobenzimidazole riboside (DRB) during ETO treatment abolished both γH2AX signal and TDP2 focus formation in G1 cells, whereas S/G2 cells showed robust γH2AX signal but no TDP2 foci (Extended Data Fig. 8e-h). TDP2 is therefore recruited, and condenses, at transcription-associated DSBs generated by TOP2β trapping.

The cAMP–PKA–CREB axis is a central transducer of both neural development and DNA damage signalling^40,52–57^. We therefore focused on PKIB, which was markedly upregulated in TDP2-mutant iPSCs, iNPCs and day 12 hCerOs (Fig. 2g-l). PKIB upregulation was accompanied by reduced phosphorylation of PKA and of its downstream effector CREB in patient-derived iPSCs, iNPCs and day 12 hCerOs (Fig. 3e-g and Extended Data Fig. 9a-d), and the same reduction occurred in TDP2 knockdown iPSCs (Extended Data Fig. 5c). Notably, iPSCs and iNPCs showed attenuated p-CREB signalling without accumulating γH2AX (Extended Data Fig. 9a-d). On transition to the organoid stage, however, sustained suppression of the cAMP–PKA–CREB axis was accompanied by marked γH2AX accumulation, indicating that the developing cerebellum is selectively vulnerable to DSBs when TDP2 is inactivated (Fig. 3h). Consistent with a widespread damage response, poly(ADP-ribose) levels were elevated in mutant hCerOs (Fig. 3j). The γH2AX signal did not colocalize with SOX2, which marks the proliferative neural progenitor pool (Fig. 3i). This spatial dissociation suggests that DSB accumulation in mutant hCerOs is uncoupled from DNA replication and is instead predominantly transcription-associated.

To test whether PKIB-mediated PKA inhibition is itself sufficient to cause the observed damage, we manipulated PKA–CREB signalling during organoid induction. The PKA inhibitor H89 reproduced the molecular signature of TDP2 inactivation in wild-type hCerOs^42,58^, causing hypoplasia and DNA damage accumulation (Fig. 3k,l and Extended Data Fig. 9f-h). This sensitivity was lineage-restricted: H89 reduced CREB phosphorylation in wild-type iPSCs without provoking DNA damage (Extended Data Fig. 9e). Conversely, the cAMP–PKA agonists cAMP and forskolin^53,55,56^ reversed the PKIB-mediated blockade in mutant hCerOs, restoring CREB phosphorylation and abolishing both the accumulated DNA damage and the hypoplasia in both mutant lines (Fig. 3m-o and Extended Data Fig. 9h,i). To ask whether PKIB dysregulation accounts fully for the mutant phenotype, we generated a PKIB-overexpressing iPSC line (Extended Data Fig. 10a). PKIB overexpression attenuated CREB phosphorylation in iPSCs but altered neither sensitivity to DNA damage nor repair kinetics (Extended Data Fig. 10b-d). During organoid induction, PKIB overexpression lowered the p-CREB/CREB ratio by raising total CREB rather than by reducing p-CREB, and produced early hypoplasia and substantial DNA damage accumulation (Fig. 3p and Extended Data Fig. 10e,f). The hypoplasia nonetheless resolved by day 12, possibly through compensatory Ca2+/CaMK and Ras–MAPK signalling^42^ (Extended Data Fig. 10g). TDP2-dependent suppression of PKIB may therefore be required to sustain the high-fidelity repair programme that cerebellar ontogeny demands. Why compensatory pathways offset PKIB overexpression yet fail to rescue the damage caused by loss of TDP2 remains to be determined.

Together these findings outline a lineage-restricted cascade in which TDP2 supports cerebellar development through the PKIB–cAMP–PKA–CREB axis. TDP2 inactivation upregulates PKIB and suppresses PKA–CREB signalling; although undifferentiated progenitors tolerate this, the same signalling collapse during cerebellar specification may compromise high-fidelity repair, leading to DSB accumulation and developmental arrest.

### TDP2 protects genome stability during early cerebellar development

To ask whether TDP2 inactivation leaves a measurable mutational footprint in vivo, we performed whole-genome sequencing (WGS) on the SCAR23 family (Fig. 4a). The homozygous patient carried more de novo mutations than the healthy and heterozygous siblings, with a pronounced excess of 1-4-base-pair (bp) deletions and a predominance of 4-bp deletions in exonic regions (Fig. 4c). Trapped TOP2 cleavage complexes generate DSBs with 4-bp cohesive ends^59,60^ (Fig. 4b), so this signature is consistent with a requirement for TDP2 in the accurate repair of TOP2-induced DSBs.

**Fig. 4.**
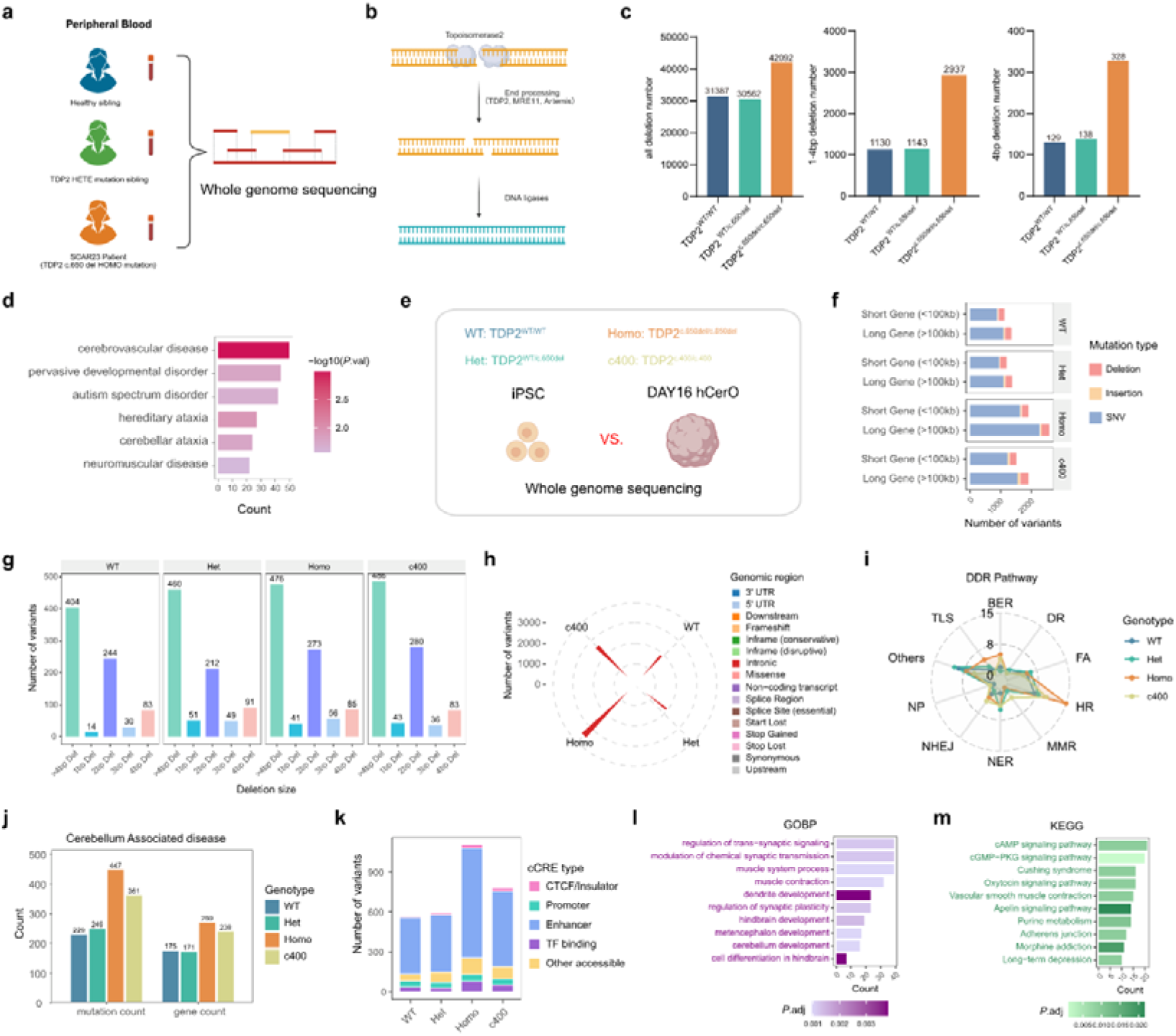
TDP2-mutant hCerOs accumulate de novo variants, including variants overlapping regulatory elements and neurodevelopmental gene panels. **a,** Design of bulk whole-genome sequencing (WGS) of peripheral blood from the SCAR23 family. **b,** Schematic of TOP2-mediated DNA cleavage and the repair of the resulting double-strand break (DSB). **c,** Numbers of all deletions, of 1-4-base-pair (bp) deletions and of 4-bp deletions in the homozygous patient and her siblings. **d,** Disease ontology enrichment of genes harbouring patient-specific 1-4-bp deletions in the homozygous TDP2 c.650del patient. **e,** Design of bulk WGS of undifferentiated iPSCs and matched D16 hCerOs from the WT, Het, Homo and c400 lines. Variants detected in D16 hCerOs but absent from the matched iPSCs are termed hCerO development-specific de novo variants. **f,** Numbers of 1-4-bp deletions and of deletions longer than 4 bp in each genotype. **g,** Distribution of deletion sizes in WT, Het, Homo and c400 hCerOs. **h,** Numbers of variants assigned to short (<100 kb) and long (>100 kb) genes, stratified by variant type. **i,** Genomic annotation of variants by SnpEff category; radial length represents the number of variants in each category. **j,** Numbers of variants in genes of curated DDR pathways. BER, base excision repair; NER, nucleotide excision repair; MMR, mismatch repair; HR, homologous recombination; NHEJ, non-homologous end joining; TLS, translesion synthesis; FA, Fanconi anaemia. **k,** Numbers of variants and affected genes in a curated panel of cerebellar development, neurodevelopmental and ataxia genes. **l,** Distribution of variants overlapping ENCODE Registry candidate cis-regulatory elements (cCREs). **m,** GO and KEGG terms for genes assigned to promoter-like or enhancer-like cCREs carrying Homo variants.

Genes harbouring patient-specific 1-4-bp deletions were enriched for biological processes associated with neurodevelopment, synapse formation and visual and sensory system development (Extended Data Fig. 11a), consistent with the visual impairment recorded in the patient. Disease ontology analysis additionally highlighted cerebrovascular disease, neurological disorders and spinocerebellar ataxia (Fig. 4d). This enrichment was specific to the homozygous patient and was not seen among genes carrying 1-4-bp deletions in either unaffected sibling (Extended Data Fig. 11b,c).

Cerebellar neurons, and Purkinje cells in particular, are more vulnerable to DNA damage during development than many other neuronal populations^5,10^. To ask whether this damage burden compromises genome stability as hCerOs develop, we performed bulk WGS on D16 hCerOs and their matched undifferentiated iPSCs for the WT, Het, Homo and c400 lines (Fig. 4e). Using a matched-sample Mutect2 workflow (Methods), we retained high-confidence variants present in D16 hCerOs but absent from the matched iPSCs, which we refer to as hCerO development-specific de novo variants. Both mutant lines acquired more deletions than the WT and Het lines, and their deletion size distributions were shifted accordingly (Fig. 4f,g). TOP2 supports transcription of the long neuronal genes that are highly expressed in neural cells, including genes for synaptic function and neuronal maturation^11,61,62^; accordingly, variants in genes longer than 100 kb, particularly single-nucleotide variants (SNVs), were more numerous in Homo hCerOs, with a similar increase in the c400 line (Fig. 4h). These findings indicate that TDP2 inactivation compromises genome stability and allows misrepair to accumulate during cerebellar development.

Most hCerO-specific variants fell within introns (Fig. 4i). To ask whether the affected genes were enriched for DDR functions, we intersected the sample-specific mutated gene sets with published DDR gene collections^63,64^. The mutant lines carried variants in more homologous recombination (HR) genes than the control lines (Fig. 4j), whereas NHEJ genes differed only modestly. TDP2-mutant hCerOs also showed a greater mutation burden in genes annotated as cerebellar-disease associated in the Human Phenotype Ontology (HPO) database^65^ than control lines (Fig. 4k). Among these, variants in ATXN2, RORA and CACNA1A were detected in both mutant lines, indicating a shared genetic vulnerability associated with TDP2 inactivation.

TOP2A and TOP2B are enriched at transcriptional regulatory elements such as enhancers and promoters, consistent with their roles in transcriptional regulation^2^, and TDP2 specifically resolves the DNA–protein crosslinks (DPCs) formed when TOP2 cleavage complexes become trapped^66^. We therefore asked whether TDP2 inactivation preferentially allows mutations to accumulate at regulatory regions, intersecting the coordinates of de novo variants with candidate cis-regulatory elements (cCREs) from the human ENCODE Registry. Homo hCerOs harboured the largest number of variants overlapping annotated cCREs, with enhancer-like elements the most represented class in all four genotypes (Fig. 4l). Because promoters and enhancers are frequent sites of DNA damage in neurons^1,2^, we performed GO and KEGG enrichment on genes whose promoter-like or enhancer-like elements carried Homo variants. These genes were enriched for cerebellum development, regulation of trans-synaptic signalling and cAMP signalling (Fig. 4m), consistent with the PKIB–PKA–CREB dysfunction identified in mutant hCerOs.

Together these findings indicate that TDP2 inactivation leads to a marked accumulation of de novo variants, particularly within cCREs, which may perturb the regulation of genes governing cerebellar development, synaptic organization and cyclic-nucleotide signalling during early hCerO differentiation.

### TDP2 inactivation redirects developmental trajectories and perturbs Purkinje cell maturation

We next examined later stages of hCerO development. By day 36 (D36) and day 63 (D63), the dominant transcriptional abnormalities had shifted to Purkinje-lineage and glial populations, which showed widespread downregulation of genes for axon guidance, synapse organization and regulation of neuron projection development (Fig. 5a). At D63, genes downregulated in Bergmann glia were enriched for cytoplasmic translation and for cilium movement involved in cell motility (Fig. 5b). This selective change in ciliary transcripts is notable because cilium-dependent signalling is required for cerebellar progenitor expansion, and loss of the ciliary protein Ttc21b in Bergmann glia perturbs glial morphology, Sonic hedgehog signalling and granule cell migration^71,72^. DEGs between Homo and Het organoids again changed in the same direction as in the Homo-versus-WT comparison (Extended Data Fig. 12a). Thus the concurrent disruption of RNA biosynthesis and early neuronal differentiation seen at D12 (Fig. 2e) is followed at later stages by a failure to sustain the transcriptional programmes required for neuronal projection and synaptic maturation.

**Fig. 5.**
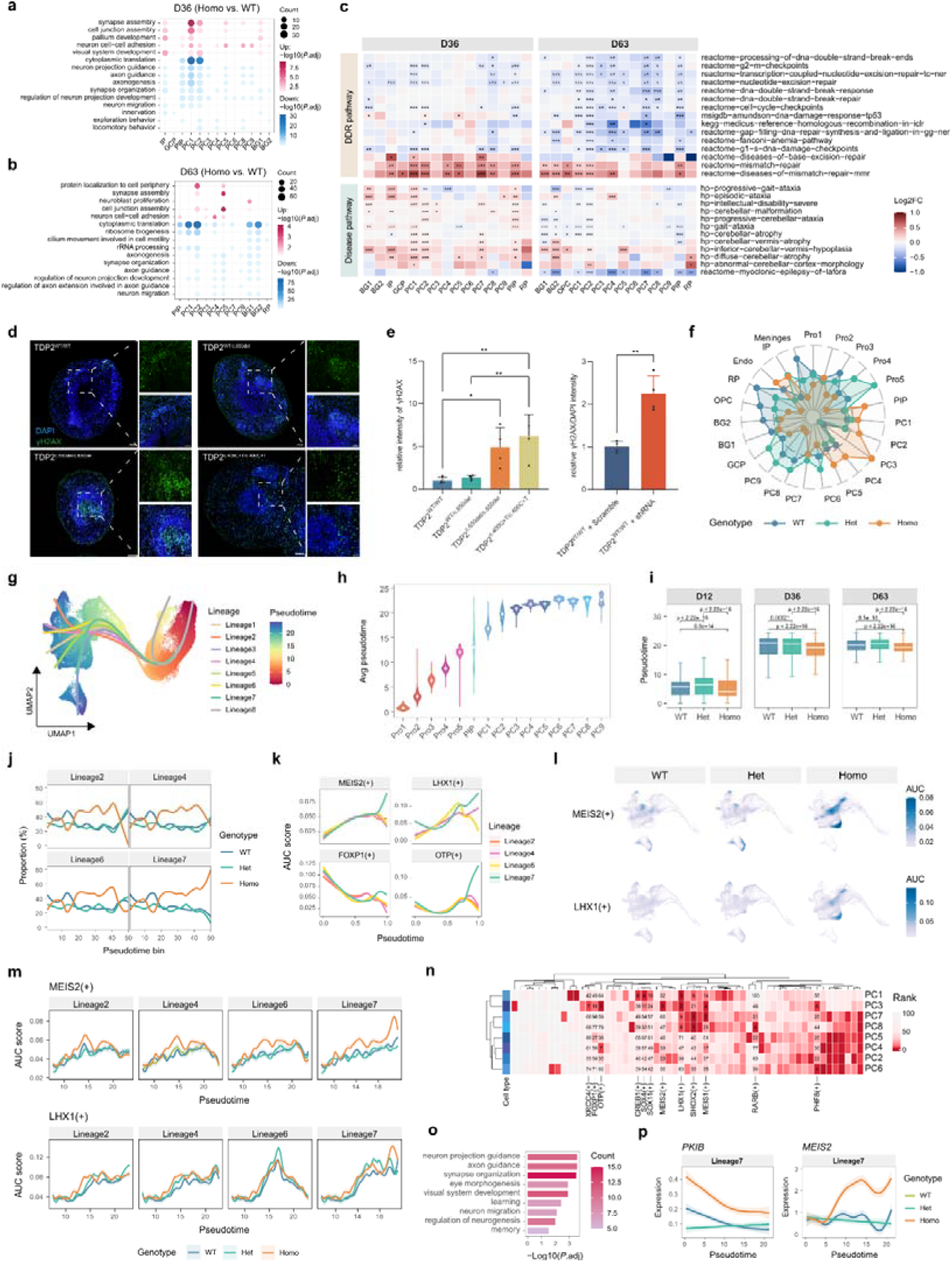
TDP2 inactivation diverts the developmental trajectories of major neuronal lineages and remodels CREB1, MEIS2 and LHX1 regulon dynamics in Homo hCerOs. **a,b,** Dot plots of enriched GO terms for DEGs in Homo versus WT hCerOs at D36 **(a)** and D63 **(b)**. **c,** Heatmaps of log2 fold change in rank-based gene set enrichment scores for DNA damage response and DNA repair (DDR) pathways (top) and cerebellum-associated disease pathways (bottom) in Homo versus WT hCerOs at D36 and D63. Red and blue denote higher and lower scores in Homo cells, respectively. Asterisks indicate Benjamini–Hochberg-adjusted Wilcoxon P values (*P.adj < 0.05, **P.adj < 0.01, ***P.adj < 0.001). **d,** Representative immunofluorescence of γH2AX in D63 hCerO sections. Scale bars, 100 μm; insets, 50 μm. **e,** γH2AX intensity in D63 hCerOs (mean intensity ± s.d., n = 4; left) and γH2AX/DAPI intensity in D63 hCerOs after shRNA-mediated TDP2 knockdown (mean intensity ± s.d., n = 4; right). **f,** Radar plot of genotype-normalized enrichment scores for each cell type in WT, Het and Homo hCerOs. **g,** Slingshot trajectories for the eight lineages identified across the major neuronal populations. **h,** Violin plots of pseudotime distributions across major neuronal populations. **i,** Box plots comparing pseudotime distributions between WT, Het and Homo cells at D12, D36 and D63. **j,** Genotype proportions along pseudotime bins for Lineages 2, 4, 6 and 7. Lineages 2, 4 and 6 were relatively enriched for WT cells, whereas Lineage 7 showed increasing representation of Homo cells towards late pseudotime. **k,** Lineage-resolved AUCell activity profiles of the MEIS2(+), LHX1(+), FOXP1(+) and OTP(+) regulons. **l,** UMAP visualizations of MEIS2(+) and LHX1(+) regulon activity stratified by genotype. **m,** Genotype-stratified pseudotime trajectories of MEIS2(+) and LHX1(+) regulon activity along Lineages 2, 4, 6 and 7. **n,** Heatmap of regulon specificity score (RSS) rankings across PC1-PC8. **o,** Enriched GO terms for genes with differential expression trajectories in both the Homo-versus-WT and Homo-versus-Het comparisons along Lineage 7. **p,** Smoothed expression trajectories of PKIB and MEIS2 along Lineage 7, stratified by genotype.

Differences in DDR pathway activity between Homo and WT cells became more cell-type dependent at D36, and by D63 many Purkinje-lineage and glial populations showed reduced activity of DSB response and repair, nucleotide excision repair, G2/M and G1/S checkpoint, and gap-filling synthesis and ligation in global-genome nucleotide excision repair (GG-NER) (Fig. 5c). Mismatch repair gene sets, by contrast, remained broadly elevated (Fig. 5c). Gene sets linked to inferior cerebellar vermis hypoplasia, cerebellar malformation, episodic ataxia and severe intellectual disability were most altered in PIP, PC1, PC2, BG1 and IP at D36 (Fig. 5c), and by D63 this broadly increased pattern gave way to a more heterogeneous and predominantly reduced profile across most subtypes (Fig. 5c). Lafora disease, an autosomal recessive progressive myoclonic epilepsy that, like TDP2-mutant SCAR23, combines epilepsy, ataxia and progressive neurodegeneration^75^, showed a related shift, its pathway score moving from widespread elevation at D12 to broad reduction at D63. Consistent with these changes, D63 mutant hCerOs accumulated more DNA damage (Fig. 5d), with a similar increase after shRNA-mediated TDP2 knockdown (Fig. 5e). The DNA damage that accumulates late in organoid development may therefore arise in part from compromised repair capacity, and the altered activity of the NER pathway raises the possibility that increased reliance on this route contributes to mutations arising from error-prone repair in mutant hCerOs.

To ask whether TDP2 inactivation altered cell-type composition during hCerO development, we calculated a genotype-normalized enrichment ratio for each cell type (Methods). WT cells were over-represented in several glial and non-neuronal populations, whereas Homo cells were enriched in Endo and IP populations and in several early PC states, most prominently PC3 (Fig. 5f). Milo analysis of local differential abundance between disease (Homo) and control (WT and Het) cells identified neighbourhoods enriched for Homo cells that were dominated by PC2, PC3 and PC4 (Extended Data Fig. 12b). TDP2 inactivation therefore changes the relative abundance of specific subtypes and cell states during hCerO development.

We next asked whether this redistribution reflected altered developmental progression within neuronal lineages (Pro1-Pro5, PIP and PC1-PC9). Slingshot identified eight trajectories across the neuronal compartment (Fig. 5g and Extended Data Fig. 12c; Methods). Average pseudotime rose from Pro1 through Pro5, increased further in PIP and was highest among PC states, supporting a progression from progenitors towards the Purkinje lineage (Fig. 5h and Extended Data Fig. 12d). The Homo-enriched PC2-PC5 states occupied earlier average pseudotime positions than the predominantly WT-enriched PC6-PC9 states. Pseudotime distributions also differed by genotype and stage (Extended Data Fig. 12e,f): Homo cells showed lower median pseudotime than WT and Het cells at D12, D36 and D63 overall (Fig. 5i), although cell-type-specific comparisons showed that this shift was not uniform across populations (Extended Data Fig. 12g). Pseudotime increased with chronological culture time in all three genotypes, but the relationship was shallowest in Homo cells, indicating slower progression along the inferred continuum (Extended Data Fig. 12h). The altered composition of Homo hCerOs is thus accompanied by preferential occupancy of intermediate Purkinje-lineage states and reduced progression to later PC states.

We found that Lineage 7 contained a higher proportion of PC3 cells in late-pseudotime bins than the other lineages (Extended Data Fig. 12i), and the proportion of Homo cells in Lineage 7 increased sharply towards its terminal pseudotime, whereas Lineages 2, 4 and 6 remained relatively enriched for WT cells (Fig. 5j and Extended Data Fig. 12j).

To examine lineage-specific gene expression dynamics during early PC differentiation, we analysed the pseudotime interval between knots 4 and 5 in each lineage (Extended Data Fig. 12k). Genes distinguishing the Homo-enriched Lineage 7 from the WT-enriched Lineages 2, 4 and 6 over this interval were associated with neuron projection guidance, neuronal migration, and learning and memory (Extended Data Fig. 12l), linking preferential occupancy of Lineage 7 to altered neurodevelopmental transcriptional programmes.

Stepwise maturation of cerebellar Purkinje cells is coordinated by lineage-specific transcription factors, and the accumulation of mutant cells in early PC states suggested that this programme was perturbed. We therefore applied SCENIC to PIP and PC populations to follow regulon dynamics through differentiation. Lineage-resolved AUC profiles revealed branch-dependent dynamics of the MEIS2(+), LHX1(+), FOXP1(+) and OTP(+) regulons, with pronounced late-pseudotime divergence between the Homo-enriched Lineage 7 and the WT-enriched Lineages 2, 4 and 6 (Fig. 5k,l and Extended Data Fig. 12m). MEIS2 encodes a TALE-class homeodomain transcription factor implicated in early neural lineage specification and neuronal differentiation^67–69^; LHX1 encodes a LIM-homeodomain factor expressed in Purkinje cell precursors that, with its paralogue LHX5, is required for their differentiation^70,71^; FOXP1 governs neuronal maturation and dendritic morphogenesis, and OTP specifies neuronal subtype identity^72–74^. Within Lineages 2, 4, 6 and 7, genotype-stratified trajectories showed that Homo cells sustained higher MEIS2(+) and LHX1(+) activity than WT cells over substantial portions of pseudotime, with the greatest divergence at late pseudotime in Lineage 7 (Fig. 5m). Regulatory divergence in Homo PC populations therefore emerges progressively during differentiation, depends on the developmental branch occupied, and involves more than a single transcription factor. At later stages, ranking by regulon specificity score (RSS) revealed substantial regulatory heterogeneity across PC1-PC8 (Fig. 5n). The RSS profile of the Homo-enriched PC3 population clustered most closely with that of PC1, an earlier PC state, rather than with more advanced PC populations, suggesting that PC3 retains an earlier-like regulatory configuration despite its assignment to the PC lineage, consistent with delayed or incomplete maturation. This abnormality persisted over chronological time: in PC3, the Homo-to-WT AUC ratios for the MEIS2(+), LHX1(+), FOXP1(+) and OTP(+) regulons all exceeded one at D36 and increased further at D63 (Extended Data Fig. 12n).

To ask whether TDP2 inactivation altered gene expression dynamics across neuronal development, we identified genes with differential trajectories between genotypes in each lineage. Focusing on the Homo-enriched Lineage 7, we intersected genes with differential trajectories in the Homo-versus-WT and Homo-versus-Het comparisons; these were enriched for neuron projection guidance, synapse organization and visual system development (Fig. 5o). Along the WT-enriched Lineages 2, 4 and 6, PKIB expression remained higher in Homo cells than in WT and Het cells across most of pseudotime, and MEIS2 showed a Homo-specific induction at intermediate pseudotime that declined towards the lineage endpoints (Extended Data Fig. 12o). Along Lineage 7, PKIB expression decreased with pseudotime but remained consistently higher in Homo cells, whereas the initial Homo-specific induction of MEIS2 was not resolved and was followed by sustained or renewed elevation at late pseudotime (Fig. 5p).

Together these findings connect gene-level expression trajectories to broader abnormalities in transcriptional regulation. Reduced CREB1(+) regulon activity in progenitors and disrupted MEIS2(+) and LHX1(+) dynamics in Purkinje cells describe a continuum of regulatory dysfunction, linking early PKIB-associated attenuation of PKA–CREB signalling in progenitors to the later mistimed activation of PC differentiation programmes. TDP2 inactivation therefore appears to create a developmental bottleneck in which mutant cells accumulate in an incompletely matured PC state.

### TDP2 inactivation disrupts cerebellar cytoarchitecture and Purkinje cell development

Purkinje cells are the principal output neurons of the cerebellar cortex, integrating input from mossy fibres, granule neurons and parallel fibres, while Bergmann glia provide specialized structural and metabolic support^40,75–78^. Purkinje cells are also more sensitive to DNA damage than other cerebellar cell types^5^, and their abnormal development causes ataxia, a disorder of balance and coordination^70^. Immunofluorescence of mature hCerOs showed reduced cell density in the molecular layer of mutant organoids, consistent with observations in Tdp2-knockout mice^7^ (Fig. 6a and Extended Data Fig. 13a). Although the broader MAP2+ neuronal population was unchanged, the density of CALB1+ Purkinje cells was markedly reduced in mutant hCerOs (Fig. 6a,b and Extended Data Fig. 13c,d). Complementation with wild-type TDP2 restored Purkinje cell numbers (Fig. 6d,e), whereas TDP2 knockdown reduced them (Fig. 6f). TDP2 inactivation therefore compromises Purkinje cell development, producing a pronounced loss of Purkinje cells in human cerebellar organoids.

**Fig. 6.**
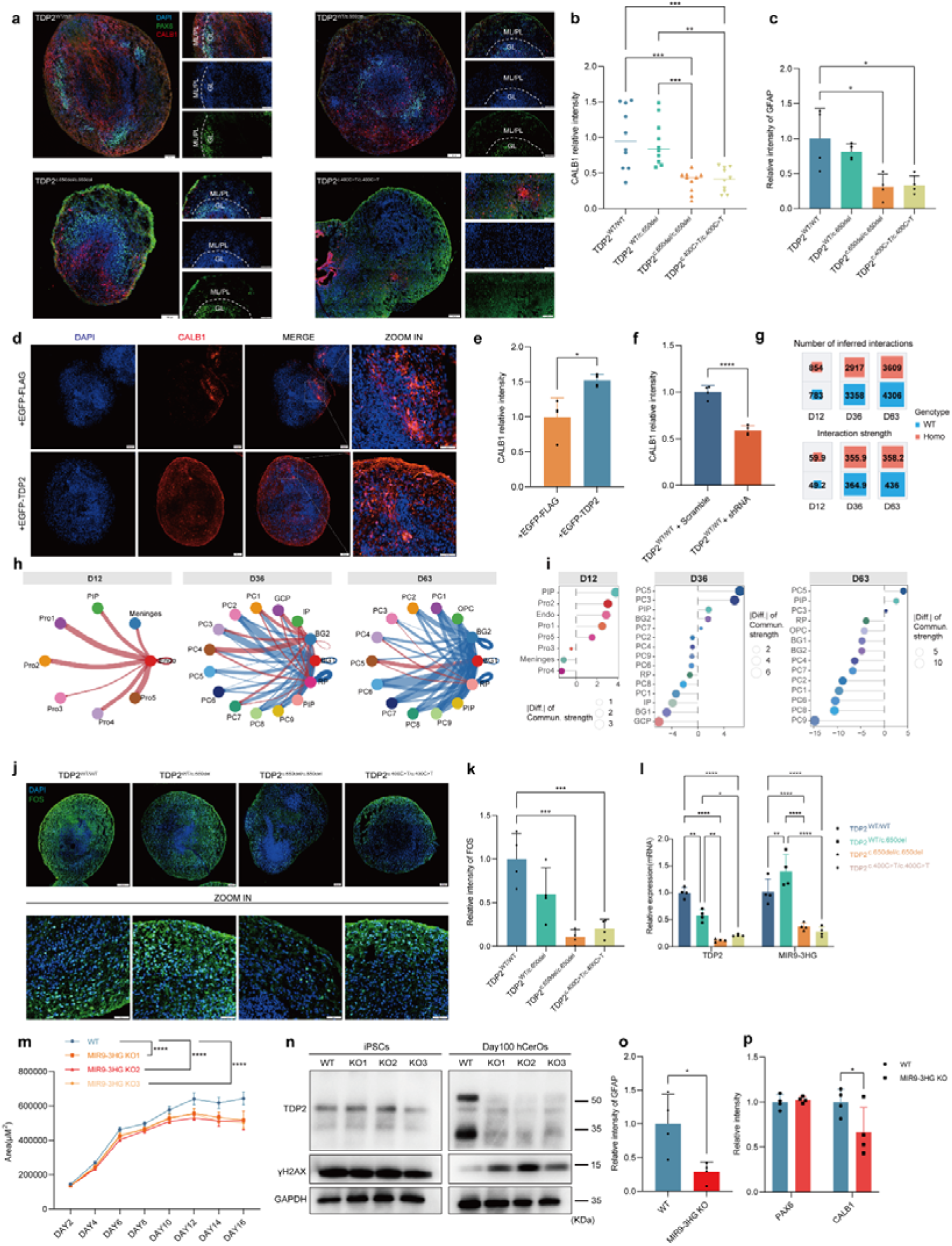
TDP2 inactivation disrupts Purkinje cell maturation and neuron–glia signalling networks. **a,** Representative immunofluorescence of day 60 hCerOs. ML, molecular layer; PL, Purkinje cell layer; GL, granule layer. Scale bars, 100 μm; insets, 50 μm. **b,** CALB1 intensity in day 60 hCerOs (mean intensity ± s.d., n = 10 organoids). **c,** GFAP intensity in day 60 TDP2-mutant hCerOs (mean intensity ± s.d., n = 4). **d,e,** Representative immunofluorescence **(d)** and CALB1 intensity **(e)** in TDP2-mutant hCerOs complemented with wild-type TDP2. Scale bars, 100 μm; insets, 50 μm (mean intensity ± s.d., n = 4). **f,** CALB1 intensity in TDP2 knockdown hCerOs (mean intensity ± s.d., n = 4). **g,h,** Numbers of inferred CellChat ligand–receptor interactions (top) and total interaction strength (bottom) in WT and Homo hCerOs at each stage **(g)**. Differential CellChat networks between non-neuronal and neuronal cells comparing Homo and WT hCerOs, from Endo at D12 and from BG1, BG2 and RP at D36 and D63; red and blue edges denote increased and decreased communication strength in Homo hCerOs, respectively **(h)**. **i,** Ranked differences in CellChat outgoing signalling strength between Homo and WT cell populations at D12, D36 and D63. **j,k,** Representative immunofluorescence **(j)** and FOS intensity **(k)** in day 60 hCerOs (mean intensity ± s.d., n = 4). Scale bars, 100 μm; insets, 50 μm. **l,** TDP2 and MIR9-3HG expression in day 100 hCerOs measured by RT–qPCR (mean ± s.d., n = 4). **m,** Cross-sectional area of MIR9-3HG-knockout hCerOs (mean ± s.d., n = 12). **n,** Representative western blots of MIR9-3HG-knockout iPSCs and day 100 hCerOs. **o,p,** Quantification of GFAP **(o)** and of PAX6 and CALB1 **(p)** in day 100 WT and MIR9-3HG-knockout hCerOs (mean ± s.d., n = 4). Statistical test: one-way ANOVA. *\*\*\*\*P*□<□0.0001, *\*\*\*P* < 0.001, *\*\*P* < 0.01, *\*P* < 0.05.

Non-neuronal populations, including choroid plexus cells, Bergmann glia and roof-plate derivatives, form niches that guide cerebellar progenitor proliferation, neuronal migration and Purkinje cell morphogenesis through morphogen, extracellular matrix and cell adhesion signalling^79^. We therefore asked whether TDP2 inactivation disrupts non-neuronal-to-neuronal communication during hCerO development. The GFAP+ Bergmann glia population was markedly depleted in mutant hCerOs (Fig. 6c and Extended Data Fig. 13c). FOS, an immediate-early gene required for neural development and function^80^, was also reduced (Fig. 6j,k), as were the immediate-early genes *FOSL1, FOSL2, NPAS4, EGR1* and *JUN*^81,82^ (Extended Data Fig. 14a,b). In mouse cerebellum after fear conditioning, Tdp2 binding was enriched at the *Fos* and *Npas4* promoters, raising the possibility that Tdp2 contributes directly to immediate-early gene transcription (Extended Data Fig. 15a-c). Disease ontology analysis of the affected genes highlighted congenital nervous system abnormalities and motor dysfunction (Extended Data Fig. 14c). CellChat analysis showed that TDP2 inactivation was accompanied by progressive remodelling of the inferred signalling network. At D12, mutant hCerOs showed more interactions and greater signalling strength than WT hCerOs; by D36 and D63 both measures were reduced, with the largest separation at D63 (Fig. 6g). These changes indicate a late decline in network connectivity across cell populations. Ranking subtypes by the Homo-minus-WT difference in summed outgoing communication strength revealed a stage-dependent shift in both the sign of the difference and the cell types contributing to it (Fig. 6i). At D12, Endo cells showed a large positive difference, and Endo-centred signalling towards Pro1-Pro5 was predominantly increased in mutant hCerOs (Fig. 6h,i), indicating strengthened choroid plexus-derived signalling from early stages. At D36, glial outputs became polarized, with a modest increase from BG2 but a marked decrease from BG1 towards Purkinje cells (Fig. 6h,i). By D63, BG1, BG2 and RP all showed negative differences (Fig. 6h,i), indicating reduced output from these non-neuronal populations. TDP2 inactivation therefore disrupts the developmental remodelling of neuron-glia communication networks in cerebellar organoids.

Many long non-coding RNAs (lncRNAs) participate in DNA repair and neurodevelopment^9,83,84^. Among them, the cerebellum-specific lncRNA MIR9-3HG, whose loss causes Purkinje cell degeneration^85^, was severely reduced in TDP2-mutant hCerOs (Fig. 6l). To test its role in cerebellar organoid development, we generated MIR9-3HG-knockout iPSC lines by CRISPR editing and differentiated them into organoids (Extended Data Fig. 16a). In mature organoids, knockout hCerOs showed the same broad suppression of immediate-early genes (*FOS, FOSB, FOSL1, NPAS4* and *EGR1*) seen in TDP2-mutant hCerOs (Extended Data Fig. 16b), and they developed pronounced hypoplasia during differentiation (Fig. 6m and Extended Data Fig. 16c,d). Unexpectedly, MIR9-3HG deletion markedly reduced expression of both the full-length and the mitochondrially localized TDP2 isoforms in mature hCerOs, while leaving TDP2 unaffected at the iPSC stage (Fig. 6n and Extended Data Fig. 16e,f). MIR9-3HG knockout also led to DNA damage accumulation during organoid development without affecting basal repair in iPSCs (Fig. 6n and Extended Data Fig. 16g), and both Purkinje cells and Bergmann glia were depleted (Fig. 6o,p and Extended Data Fig. 16h). In mouse, Tdp2 expression declined in the cerebellum with age but not in cortex or hippocampus (Extended Data Fig. 15d). TDP2 inactivation may therefore impair neuronal maturation during late cerebellar development in part through reduced MIR9-3HG expression; conversely, loss of MIR9-3HG also lowers TDP2 expression during cerebellar development and maturation.

In summary, TDP2 inactivation impairs Purkinje cell differentiation and depletes Bergmann glia. These cellular deficits disrupt intercellular connectivity and communication within the cerebellar network, stalling cerebellar development. Given that MIR9-3HG deletion likewise causes Purkinje cell and Bergmann glia loss together with DSB accumulation, the late developmental arrest and persistent genome instability that follow TDP2 inactivation may be mediated in part by reduced MIR9-3HG expression alongside altered CREB phosphorylation.

## Discussion

Our results indicate that TDP2 is required for early human cerebellar development and for genome maintenance during that period. Using iPSCs from a SCAR23 family, together with a prime-edited isogenic line of nearly identical genetic background, we found that TDP2-mutant hCerOs accumulated de novo deletions as they differentiated and repaired TOP2-trapped DSBs inefficiently. In neural cells, TDP2 was recruited both to etoposide-induced ‘junk-ended’ breaks and to clean-ended AsiSI breaks. TDP2 inactivation also impaired cerebellar organogenesis, producing atrophy. CREB couples neuronal activity to Purkinje cell differentiation and is activated under oxidative stress to support DNA repair in neural cells^39,86,87^. We find that TDP2 inactivation instead upregulates the cerebellum-enriched PKA inhibitor PKIB, suppressing PKA–CREB signalling during organoid development; as CREB phosphorylation falls, DSBs accumulate and hypoplasia follows in both TDP2-mutant and TDP2 knockdown organoids. We additionally observed dysregulated ribosome biogenesis, perturbed Purkinje cell maturation and loss of neuron–glia communication in mutant organoids. Together these findings support an essential role for TDP2 in human cerebellar development, in which its inactivation promotes mutagenesis, genome instability and cerebellar hypoplasia (Fig. 7).

**Fig. 7.**
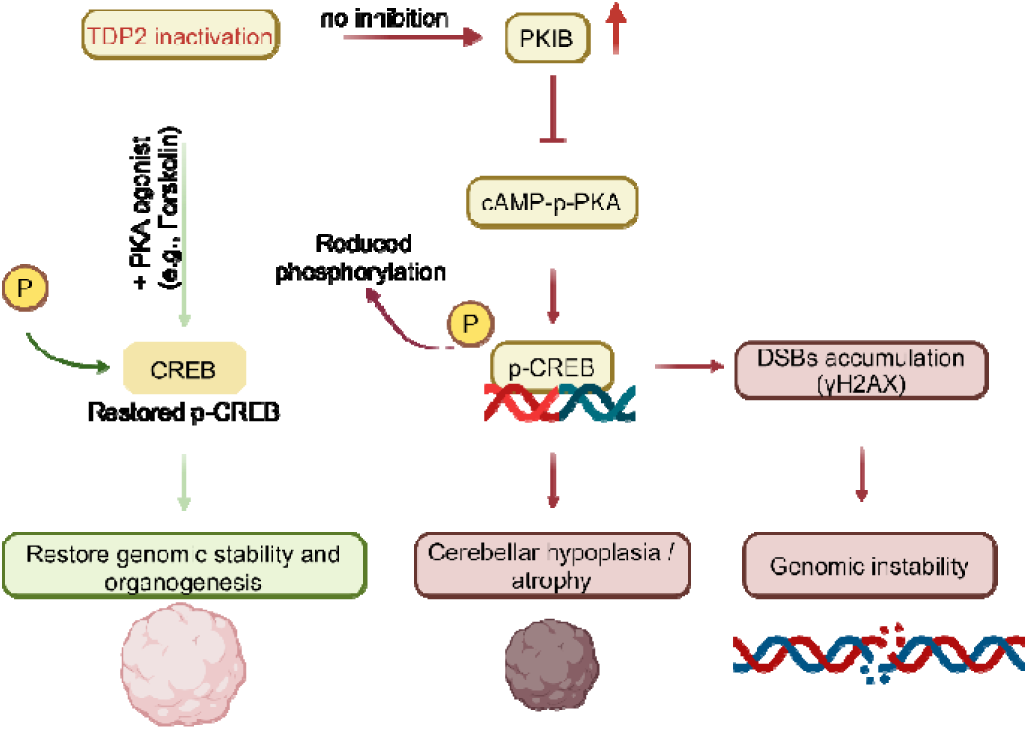
Proposed model linking TDP2 inactivation to genome instability and impaired cerebellar organoid development.

TOP2α and TOP2β relieve topological tension during transcription, replication and repair, forming a transient cleavage complex (TOP2cc) that is covalently bound to DNA and generates a DSB which is subsequently religated^88,89^. In neurons, TOP2 regulates transcription of genes required for synaptic function and plasticity^1^. Progressive cerebellar deficits are a common feature of genome instability syndromes caused by defective DNA repair^90,91^. WGS of peripheral blood from the SCAR23 family showed that TDP2 inactivation is associated with a systemic excess of 1-4-bp deletions. These correspond in size to the 4-bp cohesive ends generated by TOP2 cleavage and were enriched in genes governing neural development — particularly sensory and visual system development and synaptic function. The SCAR23 patient with TDP2 c.650del mutation had visual impairment (visual acuity 0.4 right, 0.1 left), although concurrent intellectual disability precluded fuller clinical assessment. Disease ontology enrichment of the patient’s de novo exonic 1-4-bp indels pointed to cerebellar ataxia and pervasive developmental disorders, implicating damage to cerebellar developmental genes in SCAR23 pathogenesis. To control for non-genetic contributions such as environmental exposure or infection, and to test whether misrepair is a general consequence of TDP2 inactivation, we generated a second pathogenic allele (c.400C>T) by prime editing. Differentiating both mutant lines into hCerOs showed that TDP2 inactivation intrinsically drives the accumulation of misrepair during cerebellar development. In contrast to the 1-4-bp microdeletions that dominated the family’s peripheral blood, de novo mutations in mutant hCerOs also included an excess of deletions longer than 4 bp, enriched at promoters and enhancers of neural and cerebellar development genes, whereas SNVs accumulated preferentially in long genes. Transcriptionally active neural genes thus appear particularly vulnerable to TDP2 inactivation. DNA damage continued to accumulate at late stages, but whether the genome-maintenance function of TDP2 is cell-type specific remains unresolved and will require single-cell whole-genome sequencing. Overall, our data indicate that TDP2 maintains genome stability during early cerebellar development and suggest that spinocerebellar ataxia may originate, at least in part, in DNA damage and misrepair acquired while the cerebellum is forming.

TDP2, originally described as TRAF and TNF receptor-associated protein (TTRAP), colocalizes with PML nuclear bodies and regulates rRNA biogenesis in SH-SY5Y cells^24^. Misfolded DJ-1 mutants (L166P and M26I) disrupt this nucleolar localization and sequester TDP2 into cytoplasmic aggregates when proteasome function is impaired, a mechanism implicated in sporadic and familial Parkinson’s disease and in Huntington’s disease^92^. An earlier study concluded that TDP2 does not join repair complexes at etoposide-induced DSBs in neuroblastoma cells unless the proteasome is inhibited^23,24^. We suspect that this conclusion reflected a focus on nucleolar relocalization, limited imaging resolution and the absence of co-staining for canonical damage markers such as γH2AX. Here TDP2 was recruited to, and condensed at, DSBs in a neuroblastoma line irrespective of whether the break carried a junk end (etoposide) or a clean end (AsiSI). Selective processing of clean ends by TDP2 could reduce error-prone repair in the neural genome by protecting cohesive ends from exonucleolytic attack before ligation, although why TDP2 as an end processing phosphodiesterase can be recruited to clean end is unclear and warrants further work. Notably, recruitment was preferentially triggered by TOP2β-mediated, transcription-coupled DSBs. Given that post-mitotic neurons accumulate DSBs largely through transcription, this transcription-coupled engagement positions TDP2 as an important contributor to repair fidelity in the mature nervous system. Beyond the aggregation driven by pathogenic DJ-1, we found that TDP2 mutations themselves dictate divergent aggregation behaviour: the c.650del and c.454G>A proteins assembled into aberrant condensates, whereas the c.400C>T protein lost the capacity to form foci altogether, despite near-identical predicted IDR profiles. We propose that both aberrant condensation and complete dispersion impair DNA binding and thereby abolish catalytic function. Consistent with this, the c.650del mutant showed slower repair kinetics than c.400C>T. Relative to wild-type, heterozygous and c.400C>T lines, c.650del iPSCs and iNPCs also showed downregulation of 29 C2H2-type zinc-finger genes (including PEG3, ZNF667 and ZNF736) and upregulation of the oncogenesis-associated lncRNA PCAT7, raising the possibility of a tumour-suppressive role for TDP2. Whether c.650del acts purely through loss of function, or whether its condensates impose an additional gain-of-function toxicity as has been proposed for TDP1 mutations^93^, remains an important question.

Cerebellar atrophy is a clinical hallmark of SCAR23^16,18,19,21,22^. In our model, organoid atrophy arose specifically during cerebellar lineage induction and did not occur when the cerebellum-specific inducing factor FGF8b was withheld. Transcriptomic, molecular and pharmacological analyses implicate TDP2 inactivation in the upregulation of PKIB, which is predominantly expressed in the human cerebellum. Increased PKIB suppressed phosphorylation of CREB, leading to lineage-restricted DNA damage accumulation and hypoplasia; the same changes occurred after TDP2 knockdown. Pharmacological activation of the cAMP pathway mitigated both the atrophy and the genome instability. Whether dysregulated CREB signalling is a converging mechanism across spinocerebellar ataxias require further investigation of a broader set of genotypes, including *TDP1, ATXN1, ATXN2, CACNA1A* and *ATM*.

In PKIB-overexpressing hCerOs, absolute p-CREB levels were preserved through an increase in total CREB and the activation of compensatory pathways^41^, yet the p-CREB/CREB ratio fell significantly. These organoids accumulated DNA damage and showed early atrophy, but the atrophy recovered to near-normal by about day 12. We propose that compensatory signalling, for example through calcium^41,42,53^, sustains CREB phosphorylation sufficiently for development to proceed. Why such compensation offsets PKIB elevation alone but fails in the context of TDP2 inactivation remains unresolved, as does the contribution of TDP2 functions outside the cAMP–PKA–CREB axis. Given this complexity, and although activating the cAMP–PKA cascade reduced DNA damage and partially rescued hypoplasia in hCerOs, PKA-directed monotherapy may be insufficient to reverse established neurological deficits in SCAR23, and combinatorial strategies are likely to be required. These findings nonetheless offer a framework for early intervention in spinocerebellar ataxia and associated epileptic disorders.

Loss of Tdp2 in mice reduces cerebellar molecular layer thickness and neuronal transcriptional activity without overt behavioural impairment^7,34^. In TDP2-mutant hCerOs we found reduced densities of both Purkinje cells and Bergmann glia, which would be expected to weaken inhibitory control of excitatory signalling in the cerebellum and could contribute to the ataxia and drug-resistant epilepsy seen in patients. We previously showed that loss of the lncRNA Mir9-3hg causes Purkinje cell degeneration and DNA damage accumulation in the aged mouse cerebellum^85^. Consistently, late-stage TDP2-mutant organoids showed markedly reduced MIR9-3HG expression, and MIR9-3HG-knockout organoids at the same stage recapitulated DNA damage accumulation and Purkinje cell loss. MIR9-3HG-knockout iPSCs, iNPCs and early organoids showed neither reduced CREB phosphorylation nor DNA damage, suggesting that the reduction of MIR9-3HG in TDP2-mutant organoids primarily compromises neuronal differentiation in late cerebellar development, independently of cAMP–PKA–CREB signalling. Both TDP2-mutant and MIR9-3HG-knockout organoids showed reduced FOS expression, although whether TDP2 inactivation alters neuronal activity remains to be determined by electrophysiological recording. A recent study of *TDP*2 ^fl/fl^/CamKIIa-Cre mice reported the opposite direction of effect, with increased expression of several immediate-early genes, including Fos, after Tdp2 knockout^94^. This discrepancy may reflect species- and cell-type-specific differences between the two models: that study examined forebrain excitatory neurons after conditional deletion in the postnatal brain, whereas our organoids model constitutive loss during cerebellar specification. Defining the mechanisms underlying these divergent immediate-early gene responses to TDP2 loss in mouse and human will require direct comparison in matched cell types.

Single-nucleus transcriptomics resolved how these events unfolded across cell types and stages. By D12, mutant progenitors showed broad upregulation of cytoplasmic translation, ribosome biogenesis and ribonucleoprotein complex assembly, together with increased PKIB expression and reduced CREB1 regulon activity. Because TDP2 (TTRAP) participates in ribosome biogenesis, this biosynthetic surge may represent a targeted response to nucleolar dysfunction rather than generic cellular stress. The concurrent induction of PKIB and activation of the damage response suggests a route by which progenitor stress is relayed to the differentiation programme through CREB suppression, with phenotypic consequences emerging later. Genome maintenance followed a similar temporal course: early activation of DSB response, nucleotide excision repair and checkpoint programmes gave way by D63 to reduced canonical repair activity alongside sustained elevation of mismatch repair, a pattern consistent with increasing reliance on error-prone repair. We suggest that this imbalance is self-reinforcing, because the transcriptional programmes that drive cerebellar maturation generate TOP2 cleavage complexes that cannot be resolved efficiently without TDP2, making Purkinje cell development itself a source of continuing DNA damage.

These defects were reflected in cell fate outcomes: mutant cells accumulated in early Purkinje states (PC2-PC5, particularly PC3), whereas Bergmann glia and more mature Purkinje subsets were under-represented. In the mutant-enriched Lineage 7, MEIS2 and LHX1 regulon activities, which normally decline during maturation, remained elevated at late pseudotime, with similar shifts for FOXP1 and OTP. This persistence may reflect either a failure to shut down homeobox programmes, blocking terminal maturation, or a compensatory attempt to maintain Purkinje identity in cells lacking CREB-dependent differentiation cues; regulon activity alone cannot distinguish these possibilities. Either way, the coordinated behaviour of several homeobox factors points to network-level remodelling rather than a single-factor defect. The resulting redistribution — accumulation of early Purkinje populations with depletion of inhibitory cell types — offers a cellular basis for the excitation–inhibition imbalance that may underlie ataxia and drug-resistant epilepsy in SCAR23. The non-neuronal environment was also affected: Endo-derived PTN and ADGRL signalling towards progenitors increased at D12, but by D63 outputs from Bergmann glia and roof-plate cells were broadly reduced, progressively diminishing the glial support that Purkinje cell maturation requires. Neuronal migration in the cerebellum can itself generate DSBs and may therefore contribute to the somatic mosaicism that is abundant in brain and has been linked to neuronal individuality and to neurodevelopmental and neurodegenerative disease. Whether the reduced maturation and function of Purkinje cells in mutant hCerOs is linked to damage acquired during migration remains to be determined.

Persistent DNA damage and error-prone repair in the brain are recognized contributors to ageing and to neurodegenerative disease, including Parkinson’s disease, Alzheimer’s disease and Amyotrophic lateral sclerosis. The preservation of genome stability at active regulatory regions suggests one route by which TDP2 may shield critical transcriptional programmes from damage-induced dysregulation during ageing. In summary, this study establishes an influence of DNA repair fidelity on cerebellar development and identifies a regulatory axis in which TDP2 supports genome stability and neural development through the PKIB–PKA–CREB cascade in human cerebellar organoids. Defining these developmental vulnerabilities provides a framework for understanding the aetiology of TDP2-mutant spinocerebellar ataxia and positions the cAMP–PKA axis as a candidate therapeutic target in DNA-repair-deficient ataxias, intractable epilepsies and the common neurodegenerative disorders characterized by DSB accumulation.

## Methods

### Cell lines

HEK 293T and HEK 293FT were cultured in DMEM medium (C11995500CP; Thermo Fisher Scientific) supplemented with 10% foetal bovine serum (FBS, F0193; sigma) and 1x penicillin/streptomycin (15140122; Life Technologies). SH-SY5Y cells were cultured in DMEM/F-12 (C11330500CP; Gibco) supplemented with 10% FBS and 1x penicillin/streptomycin.

### Plasmid and transfection

The shRNA sequences targeting human TDP2 were cloned into FUGW-H1-GFP-neomycin40. Human TDP2 gene was amplified from iPSCs and TDP2 c.425+1 G>A, TDP2 c.400C>T, TDP2 c.454G>A, TDP2 c.650del mutation genes were amplified from WT TDP2 iPSCs and template DNA can be removed by DpnI (R0176S; NEB) digestion following PCR. All of these genes were In-Fusion cloned in pLv-PGK plasmid with EGFP or FLAG tag using ClonExpress® MultiS One Step Cloning Kit (C113; Vazyme) as manual. AsiSI gene was synthesized by GENEWIZ company and cloned in pLV-TetON-FLAG plasmid.

HEK 293T and HEK 293FT cells were transfected with plasmid DNA with PolyJet™ DNA Transfection Reagent (SL100688; SignaGen Laboratories) following the manufacturer’s instructions. Briefly, plasmid DNA was mixed with polyjet in medium without FBS and incubated 10min at room temperature before added in cell plated. 12-18 hours after transfection, the medium was changed to a complete growth medium to reduce toxicity. The cells were collected for transfection efficiency or western blot and immunofluorescence 24-48 hours after transfection.

### Lentivirus production

To generate lentivirus, expression vectors plasmid and packaging vectors (psPAX2 and pMD2.G) were co transfected into HEK 293FT cells at the ratio of 3:2:1. The medium was changed 24 hours after transfection and medium containing lentivirus was collected 48 hours and 72 hours after transfection and ultracentrifuged at 20,000 × g for 2 h at 4 °C, the pellets were then resuspended in HBSS and stored at −80 °C until use.

### Isolation of human PBMCs

Peripheral blood samples (5–8 mL) were collected from the SCAR23 patient and healthy sibling donors in Sodium citrate-coated vacutainer tubes following informed consent. PBMCs were isolated by density gradient centrifugation using Ficoll-Paque PREMIUM, pH 5.5 - 7.5 (Cytiva, cat. no. 17-5442-02) according to the manufacturer’s instructions. Briefly, blood diluted 1:1 with PBS was carefully layered onto the Ficoll-Paque gradient and centrifuged at 400 × *g* for 10 min at 20 without brake. The mononuclear cell layer at the interphase was collected, washed twice with PBS, and either cryopreserved (PBMC medium: FBS: DMSO=7:2:1) or immediately subjected to reprogramming.

### Generation, characterization and maintenance of iPSCs

Transgene-free iPSCs were generated from PBMCs using the integration-free CytoTune-iPS 2.0 Sendai Reprogramming Kit (Thermo Fisher Scientific, cat. no. A16517) following the manufacturer’s instructions for reprogramming fibroblasts using a feeder-dependent protocol. Briefly, three days before transduction, PBMCs were plated in 24well plate for about 5 x10^5^/ml using StemPro®-34 medium (Thermo Scientific cat. 10639011, adding SCF, FLT-3, IL-3, IL-6). On day 0, CytoTune 2.0 Sendai virus was added to the cells centrifuged at 220 rpm for 90 min and incubated overnight. The medium was replaced the next day (day 1) with fresh StemPro medium. On day 2 culture dishes were prepared containing a feeder layer of colchicine pre-treatment mouse embryonic fibroblasts (MEFs). Seed the 12-well MEF culture plates with 5,000 and 25,000 live PBMCs per well in 1 mL of complete StemPro®-34 medium without the cytokines. Half-change medium with StemPro®-34 (without cytokines) on day 5. On day 7, the medium was half change with iPSCs medium (mTeSR Plus, Stem Cell Technologies cat. 100-0276). The medium was changed with iPSCs medium on day 8 and replaced the medium every day thereafter. From day 15 to 21, single iPS cell colonies were manually picked by scraping the colony and plating them in matrix (Corning cat. 354277) coating plate with iPSCs medium. Outside-family control iPS cell lines (9021 and 1011) were obtained from Coriell. All iPSCs lines were karyo typed and confirmed to be mycoplasma free. iPSCs were cultured in mTeSR Plus stem cell medium, and the medium was changed every other day. iPSCs were passaged every 4–5 days using Gentle Cell Dissociation Reagent (GCDR, STEMCELL Technologies cat. 07174) and 1/10 reseeded onto new plates coated with Matrigel (354277, Corning).

### CRISPR–Cas9 editing of iPSCs

We used Cas9 Prime editing to generate TDP2 c.400C>T mutation iPSCs from TDP2^WT/WT^ iPSCs in TDP2 c.650del SCAR23 family. On day0, single-cell suspensions from iPSCs colonies were prepared using TrypLE (Thermo Fisher Scientific cat. 12563011). On day1 (24h after plated) iPSCs clusters formed by approximately 3-5 single cells growing together and were transduced Split PEmax nCas9-BFP, Split PEmax GFP RT-MLH1 and TDP2 c.400C>T gRNA-RFP using Lipofectamine (Thermo Fisher Scientific cat. STEM00001) in Opti-MEM + CloneR2 medium (Gibco cat. 31985-062). On day2, the medium was replaced by fresh iPSCs medium. iPSCs were digested into single-cell suspension using Accutase (Stem Cell Technologies cat. 07920) on day3 and sort triple-fluorescence-positive cells using flow cytometry (BD Influx, 405, 488, 561) 1 cell per well in 96well plate. After the iPSCs colony had grown to occupy approximately 50% of the well surface area, one half of the colony was scraped using a pipette tip and collected into an Eppendorf tube for genomic DNA extraction using Rapid Animal Genomic DNA Isolation Kit (Sangon Biotech cat. B518221). Amplified PCR product containing the edit site was sent for mutation validation by Sanger sequencing. TDP2 c.400C>T mutation gRNAs, ATCTGTCAGAGAGGGCTCGA.

To generate MIR9-3HG knockout iPSCs, we transfected single-cell iPSCs (1011) cultured on MEF coating plated with MIR9-3HG KO gRNA using electrotransformation. And then, cells were selected under puromycin for 7days before packing colonies. MIR9-3HG knockout colonies were validated by qPCR.

### Generation of iPSC-induced neural progenitor cells (iNPCs)

NPCs were differentiated as previously described with minor adjustments (Robert Krencik et al., 2011). Before differentiation, iPSCs, 80% confluence on matrix coating plate in iPSCs medium, were dissociated by GCDR and reseeded on a new matrix coating plate at 1:10 to 1:15 ratio. On day0, about three days after seeding, when iPSCs hold about 30%-40% confluence, medium was replaced with NPCs medium, consisting of 1:1 DMEM/F12 and Neurobasal medium (21103-049; Gibco), 2% B27 minus vitamin a (12587001; Gibco), 1% N2 (17502001; Gibco), 2 mM GlutaMAX (35050-061; Gibco), 1x penicillin/streptomycin (15140122; Life Technologies), 2μM SB431542 (S1067; Selleck Chemicals), 3μM CHIR (S1263; Selleck Chemicals), 100μM β-mercaptoethanol (M3148; Sigma-Aldrich), 0.2μM LDN-193189 (S7507; Selleck Chemicals), 0.1mM Ascorbic acid (A4544; Sigma). On day1 to day6, NPCs medium was changed every other day. After day6, NPCs were dissociated using dispase (07923; STEMCELL Technologies) and passaged on matrix coating plate at 1:2 to 1:3 ratio. Three or four days after passage, NPCs were used for drug treatment, immunofluorescence, and RNA-seq.

### Generation of human cerebellar organoids (hCerOs)

The human cerebellar organoids were generated as Alexander Atamian et al. described previously. 80% confluent iPSC were dissociated to single cells using room temperature Accutase and were reaggregated in 96 Well Clear Round Bottom Ultra Low Attachment Microplate (Corning cat. 7007) at about 6,000 cells per well in a growth factor reduced chemically defined medium gfCDM, consisting of 1:1 IMDM (12440053; Gibco) and F-12 (11765054; Gibco), 5mg/ml >99% Bovine Serum Albumin (BSA) (I9278; Sigma), 1% chemically defined lipid concentrate (CDLC) (11905031; Gibco), 15ug/ml Apo-transferrin, and 450uM Mono-Thioglycerol (T1147; Sigma) with 7ug/ml insulin (I8830; Solabio), 10 μM of SB431542 (S1067; Selleck Chemicals), 50 ng/ml Noggin (120-10C; Peprotech), 1.7 μM CHIR (S1263; Selleck Chemicals), and 20 μM Y-27632 ROCK inhibitor (S1049; Selleck Chemicals). On Day 2, the concentration of Noggin was increased to 100 ng/mL by removing 40μL of the media and adding 50μL containing 150 ng/ml Noggin in addition to the same concentration of the previous morphogens.On day 4, 40ul of media was removed and 50ul of 200ng/mL FGF8b (C798; Novoprotein) was added to bring the final concentration in the well to 100ng/mL but without ROCKi. To promote the cerebellar cell fate, on Day 6, 40ul of media was removed and 50ul of fresh cerebellar differentiation medium (CerDM) I, containing DMEM/F-12, 20% Knockout Serum Replacement (KSR) (12660012; Gibco), 15 mg/ml Apo-Transferrin, 7 mg/ml Insulin, 2 mM GlutaMAX, 0.1 mM 2-Mercaptoethanol (M3148; Sigma), 1x penicillin/streptomycin was added to each well while supplementing the media with the same final concentration as Day4. On Day8, the final concentration of FGF8b was increased from100ng/ml to 300ng/ml and 100μl of media was added to each well after the removal of 40μl taking the final volume to 150μl. On Day10, 80μl of media was removed and supplemented with 100μl of media with the same final concentration of the drugs from Day 8. On Day 12 and 14, half media change was conducted without any drugs by removing 70μl of media and adding 75μl of fresh CerDM1.

From Day 12 to Day30, organoids were cultured in 10-cm culture plates with orbital agitation (70 r.p.m) in CerDM II, containing DMEM/F12 medium, 1% N2 (17502001; Gibco), 1% B27 (17504001; Gibco), 2 mM GlutaMAX, 1x penicillin/streptomycin and full media change was conducted every 3 days. From Day 30 to Day 60, organoids were cultured in CerDMIII, consisting of 1:1DMEM/F12 and Neurobasal medium, 1% N2, 1% B27, 2mM GlutaMAX, 5mg/ml Heparin (07980; STEMCELL), 1% Chemically Defined Lipid Concentrate (Gibco), 1x penicillin/streptomycin, 0.5 ng/ml T3 (T2877; Sigma) and 1% Matrigel (Corning cat. 354230). From day60, organoids were cultured in CerDM IV, consisting of CerDM III without Matrigel and supplemented with 14 ng/ml Brain Derived Neurotrophic Factor (BDNF) (C076; Novoprotein). Full media change was changed every 3 days on organoids after day 30.

### Cell treatments with DNA damaging drugs

#### Etoposide

HEK 293T and SH-SY5Y cells were treated with 10μM etoposide (E1383; sigma) and incubated 1 hours at 37□. iPSCs and iNPCs were treated with 1μM etoposide and incubated 1 hours at 37□.

#### Doxorubicin

HEK 293T and SH-SY5Y cells were treated with 5μM doxorubicin (D1515; sigma) and incubated 1 hours at 37□.

#### Camptothecin

SH-SY5Y cells were treated with 3μM camptothecin (CS-1049; Chemscene) and incubated 1 hours at 37□.

#### Tetracycline treatment for induced AsiSI expression

HEK 293T and SH-SY5Y cells were transfection of pTETon-FLAG-AsiSI using polyjet or lentivirus. After 48 hours after transfection, cells were treated with 0.15μg/ml - 0.2μg/ml tetracycline for 4h incubated at 37□.

### PKA pathway activation/inhibition

#### H89 treatment

For iPSCs, cells were treated with 10μM H89 (HY-15979; MedChem Express) for 1 hours at 37□. For hCerOs, 10μM or 20μM H89 was added to medium from day4 of differentiation.

#### PKA agonist treatment

iPSCs were treated with 5μM cAMP (S7858; Selleck Chemicals), 10μM cAMP or 10μM and 20μM forskolin (S2449; Selleck Chemicals) for 1h incubation at 37□. For hCerOs, organoids were treated with 10μM cAMP or 10μM forskolin from day4 of differentiation.

### G1 synchronization and transcription inhibition

To synchronize to G1 phase, SH-SY5Y cells were washed twice with serum-free medium and cultured in DMEM/F12 with 1% FBS for 72h. For transcription inhibition, cells were incubated with the 5,6-dichloro-1-β-d-ribofuranosylbenzimidazole (DRB) for 1h at 37□.

### EdU labelling of cells

HEK 293T, SH-SY5Y and iPSCs cells were incubated with 10μM EdU for 4h, and iNPCs were incubated for 8h. To harvest, cells were washed with PBS, fixed in 4% PFA for 10min, and then incubated with the click-reaction reagent for 30 min as BeyoClick EdU-488 reaction kit (Beyotime) manual described.

### RNA extraction, cDNA preparation and qPCR

Total RNA was extracted using TRIzol reagent according to the manufacturer’s instructions (Thermo Fisher Scientific). The quantity and purity of the extracted RNA were assessed with a NanoDrop 2000c spectrophotometer. Reverse transcription was performed with 1 µg of total RNA using Hifair® III 1st Strand cDNA Synthesis SuperMix for qPCR (gDNA digester plus) (Yeasen, Shanghai, China), following the manufacturer’s protocol. Quantitative PCR (qPCR) was conducted using SYBR Green qPCR Master Mix (B21203; Selleck Chemicals) on an Applied Biosystems™ QuantStudio™ 6 Flex Real-Time PCR System (Thermo fisher scientific) following the manufacturer’s instructions. Gene expression levels were normalized to GAPDH and analysed using the comparative Ct (ΔΔCt) method. Results are presented as fold change normalized to the control.

### Chromatin Immunoprecipitation (ChIP) Assay and ChIP-qPCR

Chromatin immunoprecipitation was performed using a ChIP assay kit (cell signalling technology, #9005) as described by the manufacturer. Briefly, mouse tissue (about 200mg) was crosslinked by treatment with formaldehyde (1% final). Then nuclei and chromatin were digested using Micrococcal Nuclease and ultrasonication. Sheared chromatin was immunoprecipitated variously with antibodies against IgG, γH2AX, TDP2, and using ChIP-Grade Protein G Magnetic Beads. Crosslinked proteins were then eluted from the beads in a solution containing NaCl and Proteinase K. Crosslinks were reversed by incubation at 65°C overnight, following which DNA was purified using spin columns. For ChIP-qPCR, the enrichment fold was calculated relative to the IgG ChIP sample and further normalized to control genomic regions with constitutively open or closed chromatin.

### Western blotting

Pre-cold RIPA buffer (P0013B;Beyotime) with Protease Inhibitor Cocktail (EDTA-Free) (HY-K0010; MedChem Express) and phosphatase inhibitor cocktail (B15001; Selleck) was used to lyse cerebral organoids or cells for 30 min on ice, followed by sonication using a Fisherbrand™ Model 705 for 10% amplification, 5 s on and 10 s off, 5 cycles. Protein concentration was measured using Pierce BCA Protein Assay Kit (23227; Thermo Fisher Scientific) at an absorbance wavelength of 562 nm. Equal amounts of protein were resolved on SDS–PAGE (Bio-Rad) gels before transfer to Immobilon-P, PVDF membrane (IPVH00010; Merck). Membranes were blocked with 5% BSA (ST023; Beyotime) in TBS-Tween-20 (TBS-T) for 1 h at room temperature before incubating with primary antibodies overnight at 4 °C and secondary antibodies overnight at 4 °C and 1 hour at room temperature, respectively.

### TDP2 enzyme activity assay

The enzyme activity of TDP2 mutation protein was measured by TDP2 Assay Kits (170524; Inspiralis). Briefly, Cells were lysed and whole cells protein were extracted as described. Equal amounts of protein were supplied in Dilution buffer. And then, protein sample was mixed with substrate and volume was added up to 20μl by Assay buffer, after which the mixture was incubated 30min at 37 before measured fluorescent signal at 520nm using ELISA at 37 (recorded for 1 hours at rate of 1min per measurement).

### Cells staining

iPSCs, NPCs, HEK 293T and SH-SY5Y cells cultured on glass coverslips were fixed in 4% PFA for 10min at room temperature and blocked in blocking media (5% normal goat serum, 0.2% Triton X-100 in PBS) for 1h at room temperature. Cells were incubated with primary and secondary antibodies diluted in antibody dilution media (1% normal goat serum, 0.1% Triton X-100 in PBS) overnight at 4 °C and 1h at room temperature, respectively, followed by washing four times with PBS. Cells were incubated with DAPI (HY-D0814; MedChem Express) for 10min at room temperature after which coverslips were mounted onto glass slides with Fluoromount-G (0100-01; SouthernBiotech). Images were taken with Leica SP8 confocal microscope.

### Organoids frozen section and immunofluorescence

Organoids were fixed in 4% paraformaldehyde for 1h at room temperature. Washing with PBS for three times, organoids were incubated in10% - 30% sucrose solution at 4 for gradient dehydration. Organoids were then embedded in Tissue-Tek O.C.T. compound (62550; Sakura) and cryosectioned by Leica CM3050 S at 15μm thickness onto glass slides. Slides were incubated at room temperature overnight after which slides washed 3x with PBS before a 1-hour incubation in blocking media (described as above). After permeabilization and blocking, immunofluorescence was done as described above. Images were taken with Olympus SpinSR confocal microscope.

### Live-cell imaging

After cells were transfected with EGFP tag plasmid by lentivirus, cells were incubated with Hoechst in PBS for 10min at 37□. Cells were washed with PBS for three times and medium was replaced with DMEM or DMEM/F12 without phenol red. Cells were imaged for 1 hour at rate of 5min per frame using the HIS-SIM. Images were processed and analysed using ImageJ.

### TDP2 mutant protein structure and IDR prediction

Structures of wild-type and mutant TDP2 were predicted with AlphaFold 3 and rendered with per-residue model confidence (pLDDT). Intrinsic disorder was scored per residue with IUPred2A, and a residue was considered disordered if its IUPred score exceeded 0.5.

### RNA-seq

RNA-seq was performed by GENEWIZ using an Illumina Novaseq platform. Read quality was assessed using FastQC (v0.10.1) to identify sequencing cycles with low average quality, adapter contamination, or repetitive sequences. Filter Fastq files were trimmed and aligned to the human reference genome (hg38) using STAR (v.2.7.11a). The mapped reads were assembled with FeatureCounts (v.2.0.8), and differential gene expression was analysed using DESeq2 (v.1.42.1) based on read counts. Gene Ontology analysis of differentially expressed genes with gene symbols was performed using MetaScape.

#### Whole-genome sequencing (WGS)

For each sample, >200μg genomic DNA was randomly fragmented by Covaris to an average size of 300-350bp. The fragments were then treated with End Prep Enzyme Mix for end repairing, 5’ Phosphorylation and 3’ adenylated, to add adaptors to both ends. Size selection of the adaptor-ligated DNA was performed by DNA Cleanup beads. Each sample was then amplified with P5 and P7 primers carrying sequences which can anneal with flowcell to perform bridge PCR and index allowing for multiplexing. The PCR products were cleaned up and the final library was qualified. The qualified libraries were sequenced pair end 150bp on the illumina Hiseq Xten/Novaseq / MGI2000 System. Perform a preliminary statistical analysis of the raw data. Then, optimize the sequencing raw data by using Cutadapt software (version 1.9.1), remove the primers and adapter, the bases with the quality score below 20 at both ends, and the reads with N base ratio more than 10%. Finally, statistical analysis of clean data after QC. Clean Data are compared with reference genomes by Sentieon (or BWA), sequenced, and regenerated into BAM files. The Sentieon software(or GATK haplotypecaller)^95^ were used to call Germline SNV/InDel and the variants were annotated by Annovar^96^. Somatic variation analysis was performed by Sentieon(or Mutect) Somatic Pipeline when necessary.

### Single nucleus isolation

Nucleus was isolated using Shbio Nuclei Isolation Kit (SHBIO,#52009-10, China). Nuclei were counted with a cell counter (Thermo Fisher, America). Using a Chromium Single Cell 3′ Library and Gel Bead Kit v3 (10X Genomics),nuclei were immediately loaded onto a Chromium Single Cell Processor(10X Genomics) for barcoding of RNA from single nuclei.

## 10× library preparation and sequencing

Beads with unique molecular identifier (UMI) and cell barcodes were loaded close to saturation, so that each cell was paired with a bead in a Gel Beads-in-emulsion(GEM). After exposure to cell lysis buffer, polyadenylated RNA molecules hybridized to the beads. Beads were retrieved into a single tube for reverse transcription. On cDNA synthesis, each cDNA molecule was tagged on the 5’end (that is, the 3’end of a messenger RNA transcript) with UMI and cell label indicating its cell of origin. Briefly, 10× beads that were then subject to second-strand cDNA synthesis, adaptor ligation, and universal amplification.Sequencing libraries were prepared using randomly interrupted whole-transcriptome amplification products to enrich the 3’ end of the transcripts linked with the cell barcode and UMI. All the remaining procedures including the library construction were performed according to the standard manufacturer’s protocol (CG000204 Rev B). Sequencing libraries were quantified using a High Sensitivity DNA Chip (Agilent) on a Bioanalyzer 2100 and the Qubit High Sensitivity DNA Assay (Thermo Fisher Scientific). The libraries were sequenced on NovaSeq6000 (Illumina) using 2×150 chemistry.

### snRNA-seq data analysis

Raw FASTQ files were processed using Cell Ranger (v7.0.1, 10x Genomics)^97^ with default parameters to align reads to the human reference genome (GRCh38-1.2.0). Ambient RNA contamination was estimated and removed from the unfiltered matrix using SoupX (v1.6.2)^98^. The correctedcount matrices were further analysed using Seurat (v5.1.0)^99^. Quality control was performed on individual libraries prior to data integration by removing nuclei that failed to meet established thresholds for detected genes, total unique molecular identifiers (UMIs), and mitochondrial transcript fractions. Putative doublets were identified and removed using DoubletFinder (v2.0.4)^100^. Data were log-normalized, and the 2,000 most variable features were selected for dimensionality reduction. Batch effects across samples were corrected using the fast mutual nearest-neighbour (fastMNN) method^101^. The first 30 dimensions of the principal component analysis (PCA) embedding were used to construct a k-nearest-neighbour graph, perform graph-based clustering and generate uniform manifold approximation and projection (UMAP) visualizations.

### Cell type and subtype annotation

Major cell types and regional identities were annotated by combining established marker genes (e.g., *MKI67*, *TOP2A*), top marker genes identified by Seurat’s FindAllMarkers function, and reference-based cell type annotation mapping via Seurat’s label transfer function. Specifically, regional annotations were transferred from published snRNA-sequencing data of human neural organoids^38^, while major cell classes were mapped using human cerebellar organoid reference single-cell RNA-seq data^36^. Subclustering of major populations yielded 24 transcriptionally distinct cellular subtypes. The final annotated states encompassed progenitor cells (Pro1–Pro5), endothelial cells (Endo), interneuron progenitors (IP, and PIP, PAX2+ interneuron progenitors), Purkinje cells (PC1-PC9), Bergmann glia (BG1/BG2), granule-cell progenitors (GCP), meningeal cells, oligodendrocyte progenitor cells (OPC), roof-plate cells (RP), and other non-neuronal populations. Cellular subtypes were retained across all downstream analyses.

### Differential gene expression analysis

To identify differentially expressed genes (DEGs) among genotypes, the Seurat’s function FindMarkers with MAST algorithm was used with the thresholds log2 fold change□>□0.2, adjusted P-value□<□0.05 and min.pct□>□0.1.

### Cell-cell communication analysis

Ligand-receptor communication was inferred using the R package CellChat (version 2.1.2)^102^. Differential CellChat analysis was performed to compare inferred cell–cell communication networks between Homo and WT human cerebellar organoids (hCerOs). The total outgoing communication strength of selected pathways for each cell type as a sender was calculated by summing the inferred communication probabilities across all recipient cell types and significant ligand–receptor interactions. Differences in outgoing signalling communication strength (Homo minus WT) were then calculated to identify cell types with altered signalling outputs.

### Cellular composition enrichment and differential abundance analysis

Cellular subtype composition representation was summarized using a genotype-normalized enrichment ratio. For cell type and genotype, the ratio was calculated as, where is the number of nuclei of genotype assigned to cell type, is the total number of nuclei of genotype, is the total number of nuclei assigned to cell type, and is the total number of nuclei in the analysis. Ratios greater than one indicate representation above that expected from the overall frequency of the genotype. Local differential abundance was assessed using MiloR (v2.0.0)^103^ for the comparison of cellular composition between disease (Homo cells) and healthy control (WT and Het cells) for all cell types. Neighbourhoods were coloured by the log-transformed ratio (log2 fold change) of cell abundance between the disease and healthy control groups for visualization.

### Developmental-potential estimation

Developmental potential was estimated using CytoTRACE2 (v1.1.0)^104^ with the count matrix of neuronal cells, comprising subtypes Pro1-Pro5, PIP and PC1-PC9. The analysis returned a continuous potency score and categorical potency assignments, which were subsequently transferred to the integrated Seurat object for UMAP visualization.

### RNA-velocity analysis

Spliced and unspliced transcript counts were generated using velocyto (v0.17.17)^105^ and matched to the filtered and normalized expression matrices of neuronal cells, comprising Pro1-Pro5, PIP and PC1-PC9. RNA velocity was estimated with scVelo (v0.3.3)^106^. Velocities were inferred using the stochastic model, and the velocity graph was visualized onto the integrated UMAP coordinates calculated in Seurat pipeline. Streamlines were displayed separately for WT, Het and Homo groups.

### Lineage reconstruction and pseudotime analysis

Developmental trajectories of the major neuronal cells, comprising Pro1–Pro5, PIP, and PC1-PC9 were reconstructed using the Slingshot (v2.7.0) R package^107^. Pseudotime was computed by applying a simultaneous principal curves algorithm to construct smooth curves from the minimum spanning tree. Based on the lineages inferred by Slingshot, generalized additive models (GAMs) were fitted using the fitGAM function with nknots = 6 via the tradeSeq (v1.18.0) R package to model gene expression dynamics along pseudotime^108^. Subsequently, lineage- and genotype-specific expression patterns were evaluated using tradeSeq functions: patternTest was used to identify genes with significant expression changes across lineages; earlyDETest was used to detect differential expression between lineages at early stages of PC differentiation; and conditionTest was used to test for differences in expression trajectories between distinct genotypes.

### Gene-regulatory-network inference and regulon analysis

Gene-regulatory networks (GRNs) of neuronal cells (Pro1-Pro5, PIP and PC1-PC9) were inferred using the pySCENIC (v0.12.1) workflow^109^. Gene co-expression modules were first identified with GRNBoost2. Putative regulons were then refined by cis-regulatory motif enrichment analysis, and regulon activity was quantified for each nucleus as an area-under-the-recovery-curve (AUCell) score. The pySCENIC-derived AUCell matrix was subsequently imported into R for downstream analyses. Regulon specificity scores (RSSs) were calculated for each cellular subtype using R package SCENIC (v1.3.1). Regulons were ranked independently across PC1–PC8 according to their RSSs, and the union of the ten highest-ranking regulons from each PC state was visualized. In PC3, genotype divergence was summarized as the Homo-to-WT ratio of the mean AUCell score at D36 and D63 for MEIS2(+), LHX1(+), FOXP1(+) and OTP(+). Regulon activity was modelled along Slingshot pseudotime separately for each lineage and genotype using generalized additive model.

### Whole-genome sequencing analysis

Bulk whole-genome sequencing (WGS) was performed on undifferentiated iPSCs and matched D16 hCerOs from the WT, Het, Homo and c400 lines. Candidate hCerO-specific variants were called using a matched-sample Mutect2 workflow via GATK (v4.6.2.0). Only variants marked PASS and detected in D16 hCerOs but not in the matched iPSC sample were retained for downstream analysis. PASS variants were annotated with SnpEff (v5.1d). Variant-associated genes longer than 100 kb were defined as long genes and those shorter than 100 kb as short genes. To assess overlap with regulatory DNA, hCerOs-development specific newly formed variants were intersected with the human ENCODE Registry of candidate cis-regulatory elements (cCREs). Each overlapping variant was assigned to its annotated cCRE class, including promoter-like, enhancer-like, CTCF/insulator, transcription-factor-binding or other accessible elements.

### Functional enrichment analysis

Gene Ontology over-representation (GO) analysis was performed using the clusterProfiler R package (v4.2.1)^110^.Terms with adjusted-P > 0.05 were filtered. Single-cell gene-set activity was quantified using the AUCell method in the irGSEA R package (v3.3.4)^111^. Pathway acitivity scores were compared between genotypes using a two-sided Wilcoxon rank-sum test.

### Statistical analysis

Statistical analyses of snRNA-seq data were performed in R (v4.4.1). Comparisons of pathway activity scores between genotypes used two-sided Wilcoxon rank-sum tests with Benjamini–Hochberg correction for multiple testing; adjusted P values are reported as P.adj. All other experiments were analysed in GraphPad Prism (version 9.0.0). For each experiment, n denotes the number of independent biological replicates, or the number of organoids where stated in the figure legend; the unit of replication is defined in each legend. Data are presented as mean ± s.d. unless otherwise indicated. Comparisons of more than two groups used one-way or two-way ANOVA as specified in each legend, followed by the stated post hoc test with correction for multiple comparisons. Normality and equality of variance were assessed before parametric testing. All tests were two-sided and P < 0.05 was considered significant. No statistical method was used to predetermine sample size. Experiments were not randomized, and investigators were not blinded to genotype during data collection and analysis, except where stated. No data were excluded from the analyses.

## Data visualization

Data visualization was performed in R using ggplot2 (version 3.4.2) package. Schematic diagrams were created with BioRender (BioRender.com).

## Data availability

The snRNA-seq, bulk RNA-seq and whole-genome sequencing data generated in this study have been deposited in the NCBI (PRJNA1553986) and the data will be made publicly available upon publication. Human whole-genome sequencing data from the SCAR23 family are subject to participant-consent restrictions and are available from the corresponding author on reasonable request under a data transfer agreement; the access procedure and expected response time are given in the Supplementary Information. Published datasets used in this study comprise the human cerebellar organoid scRNA-seq reference (ref. 36), the human neural organoid snRNA-seq reference (ref. 38), the ENCODE Registry of candidate cis-regulatory elements and the curated DDR gene sets of refs 63,64. Source data for all graphs are provided with this paper.

## Code availability

All custom code generated in this study is available at https://github.com/mingmg99/hCerOs and will be made publicly available upon publication. Software versions used for all analyses are given in the relevant Methods sections.

## Ethics statement

All human studies were approved by the Institutional Review Board of Department of Neurology, the Seventh Affiliated Hospital, Sun Yat-sen University, Shenzhen, Guangdong Province, China 518107 (approval number KY-2024-046-01) and were conducted in accordance with the Declaration of Helsinki. Written informed consent was obtained from all participants in the SCAR23 family, and from the parents or legal guardians of participants unable to provide consent in their own right, for the collection of peripheral blood, the derivation and use of induced pluripotent stem cell lines, genome sequencing, and the publication of clinical and imaging data. Consent for publication of the magnetic resonance images shown in Extended Data Fig. 1a was obtained separately. Derivation and use of human iPSC lines were approved by the stem cell research oversight committee of Interdisciplinary Research Center on Biology and Chemistry (IRCBC), Shanghai Institute of Organic Chemistry, Chinese Academy of Sciences. The commercially obtained control iPSC lines were used under the terms of their original consent. All mouse experiments were approved by the Institutional Animal Care and Use Committee of Interdisciplinary Research Center on Biology and Chemistry (IRCBC), Shanghai Institute of Organic Chemistry, Chinese Academy of Sciences (202003080014) and complied with relevant national guidelines; the strain, sex, age and housing conditions of the animals used are given in the Methods.

## Reporting summary

Further information on research design is available in the Nature Portfolio Reporting Summary linked to this paper.

## Acknowledgements

We thank the members of the SCAR23 family for their participation. We thank the staff members of the Integrated Laser Microscopy System (https://cstr.cn/31129.02.NFPS.CLMIS) at the National Facility for Protein Science in Shanghai (https://cstr.cn/31129.02.NFPS) for providing technical support and assistance in data collection and analysis. We also thank Dr. Xue qi Gong of Fudan University and Dr. Lu Zhang of IRCBC for kindly helping with experiments, Dr. Bo yuan Yao of Department of Neurosurgery, Huashan Hospital for helping with RNAseq analysis.

## Author contributions

W.P.W. performed the majority of the experiments. G.M.M. performed WGS and snRNAseq data analyses. F.W. and S.Y.D. performed wt and Het iPSC lines establishment. Y.Q.Z. performed SCAR23 clinical data collection. M.M.L. performed PBMCs isolation.

## Additional information

Supplementary Information is available for this paper. Correspondence and requests for materials should be addressed to L.M.Z., Y.D.S. or W.Y.W. Peer review information and reprints and permissions information will be supplied by the publisher.

## Extended Data

**Extended Data Fig. 1.**
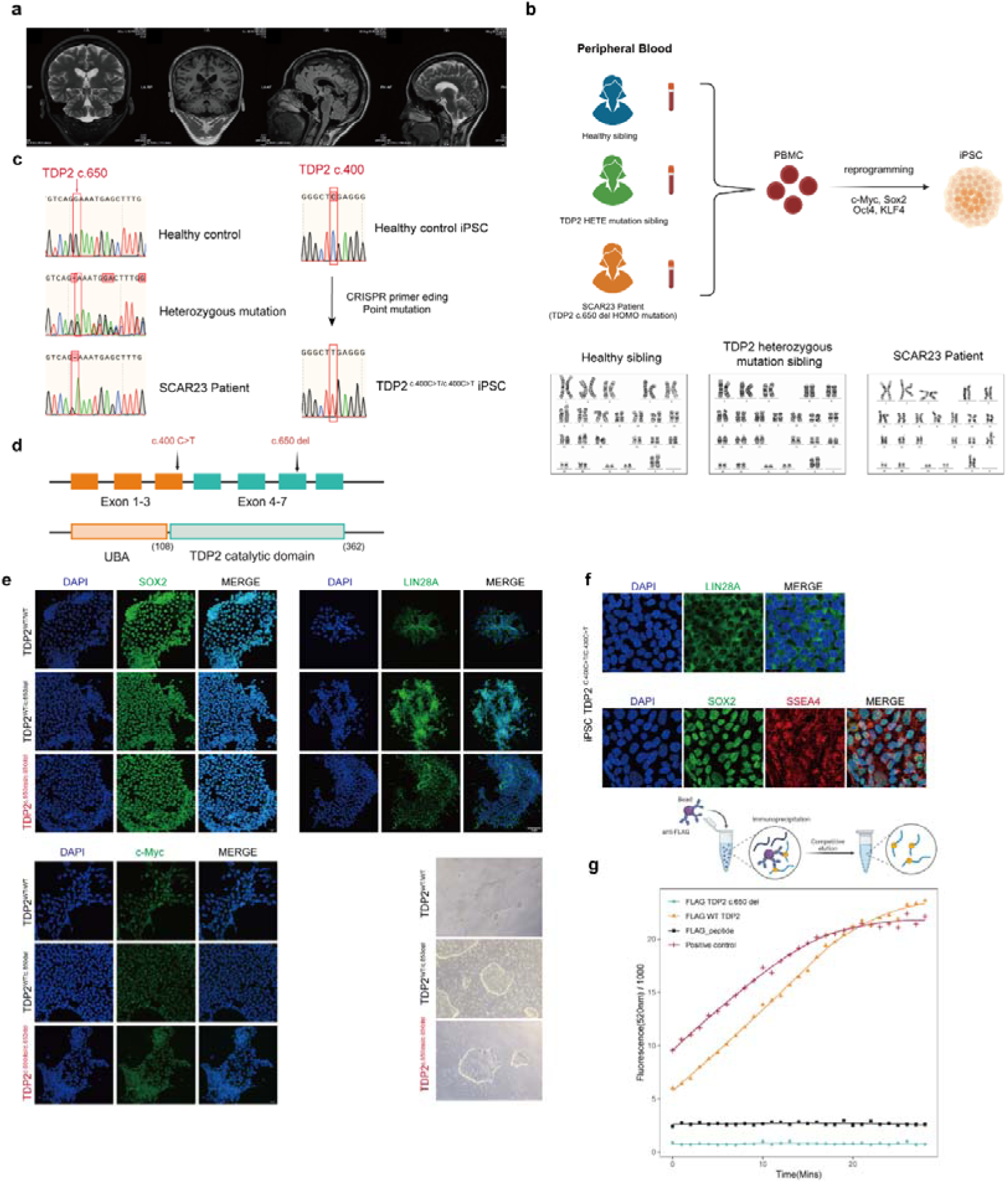
Design and generation and characterization of iPSC cells. **a,** Magnetic Resonance Imaging (MRI) of SCAR23 patient with the homozygous TDP2 c.650del mutation. **b,** Schematic of the workflow for iPSC reprogramming and Karyotypes for healthy sibling (wlid-type TDP2), TDP2 heterozygous mutant sibling (TDP2^WT/c.650del^), and SCAR23 patient (TDP2^c.650del/c.650del^). **c,** Sanger sequencing validation delineating the patient-derived and prime-edited (*TDP2* c.400C>T) isogenic mutant lines from healthy sample in SCAR23 family. **d**, Schematic representation of the *TDP2* locus mapping the relative positions of the c.400C>T and c.650del mutations across the structural domains. **e,f,** Representative immunostaining of iPSCs for pluripotency markers SOX2, LIN28A, and c-Myc. Bright-field images showed cloning morphology. Scale bars, 10 μm **(e)**. Scale bars, 23.2 μm **(f)**. **g,** In vitro biochemical tracking of purified TDP2 protein kinetics establishes the complete catalytic ablation of phosphodiesterase activity in the SCAR23-associated TDP2 c.650del mutant variant.

**Extended Data Fig. 2.**
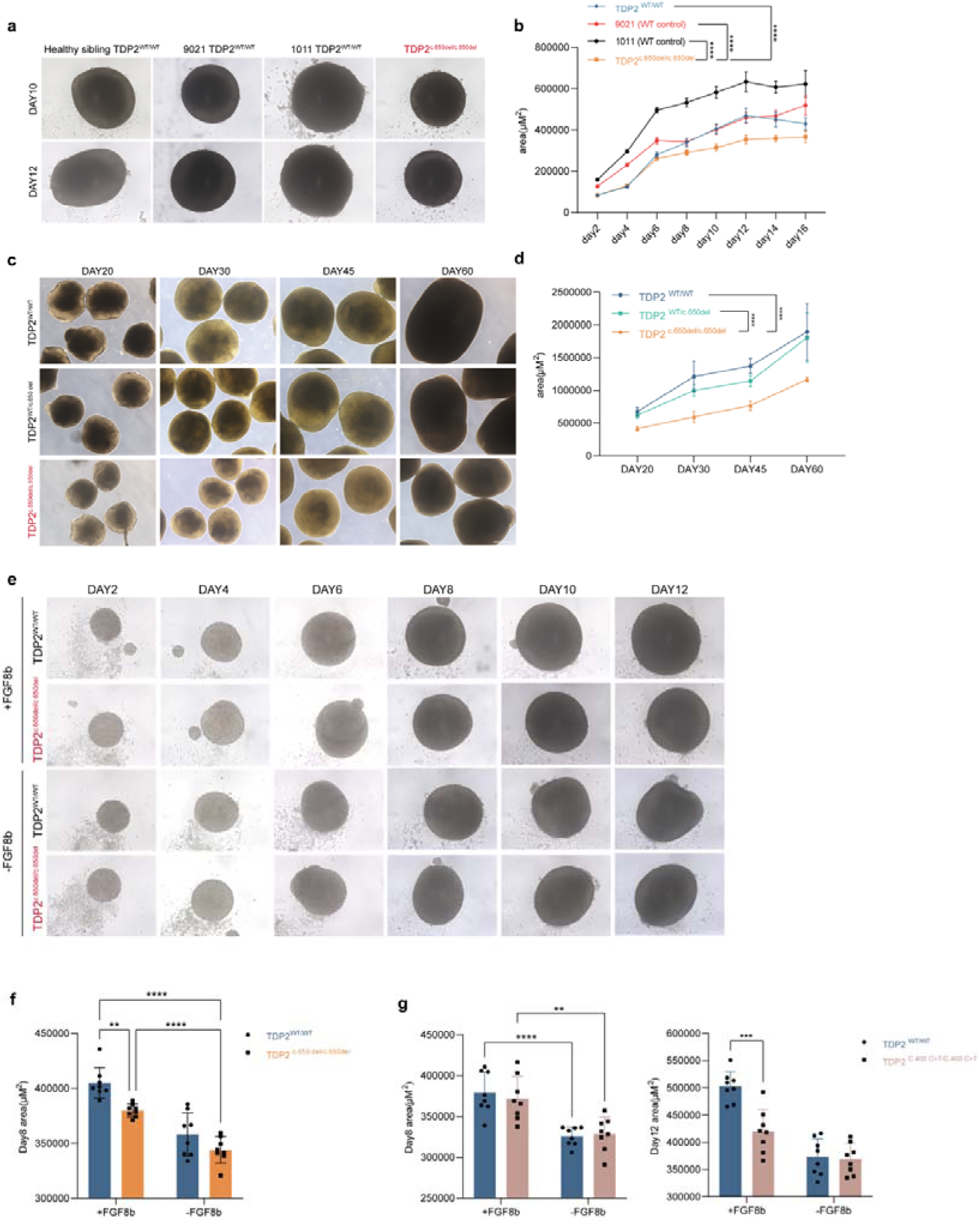
Robust validation of lineage-specific cerebellar atrophy across independent TDP2-mutant models. **a,b,** Representative brightfield of day 10 and day 12 hCerOs derived from WT TDP2, TDP2 c.650del mutant lines, and healthy individuals from outside the SCAR23 family **(a)**. Quantitative analysis **(b)** of hCerOs area from day 2 to day 12 induction (mean ± SD, n = 12). Scale bars, 50 μm. **c,d,** Representative brightfield and quantitative analysis of hCerOs area from day 20 to day 60 induction (mean ± SD, n = 6 for day 20 and day 30, n = 5 for day 45, n = 4 for day 60). Scale bars, 400 μm. **e,** Representative brightfield of hCerOs area of standard and FGF8b-deprived conditions in WT TDP2, TDP2 c.650del and TDP2 c.400C>T mutant lines. Scale bars, 200 μm. **f,g,** Quantitative analysis of standard and FGF8b-deprived conditions day 8 TDP2 c.650del hCerOs area **(f)** and day 8-12 TDP2 c.400C>T hCerOs **(g)** compared with WT TDP2 hCerOs (mean ± SD, n = 8). Statistical test: two-way ANOVA. *\*\*\*\*P* < 0.0001, *\*\*\*P* < 0.001, *\*\*P* < 0.01, *\*P* < 0.05.

**Extended Data Fig. 3.**
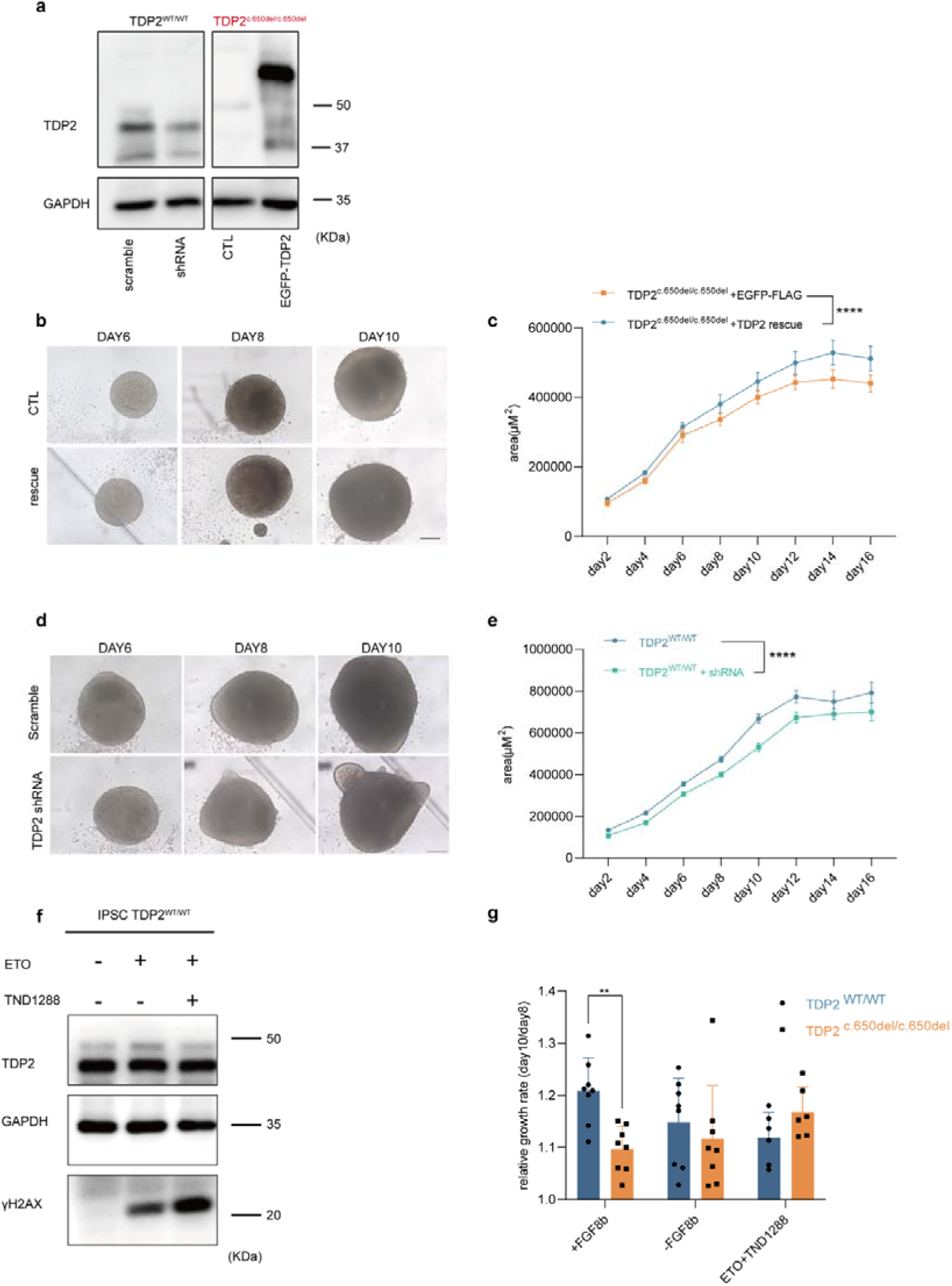
Comparison of hCerO area of WT TDP2-knockdown and TDP2 c.650del rescue line. **a,** Representative western blot of TDP2 expression in WT TDP2 line knockdown via shRNA and TDP2 c.650del mutant line rescue via wild-type TDP2 overexpression. **b,c,** Representative brightfield images and quantitative analysis of hCerOs area of wild-type TDP2 rescue hCerOs in day 6 to day 10 induction. **d,e,** Representative brightfield images and quantitative analysis of hCerOs area of TDP2 knockdown hCerOs in day 6 to day 10 induction. **f,** Representative western blot of TDP2 and γH2AX expression in TDP2 inhibition WT iPSCs by TND1288 and etoposide. **g,** Quantitative analysis of relative growth rate (day 10/day 8) for standard, FGF8b-deprived and TDP2 inhibition conditions in WT and TDP2 c.650del hCerOs (mean ± SD, n = 8). Statistical test: two-way ANOVA. *\*\*\*\*P* < 0.0001, *\*\*\*P* < 0.001, *\*\*P* < 0.01, *\*P* < 0.05.

**Extended Data Fig. 4.**
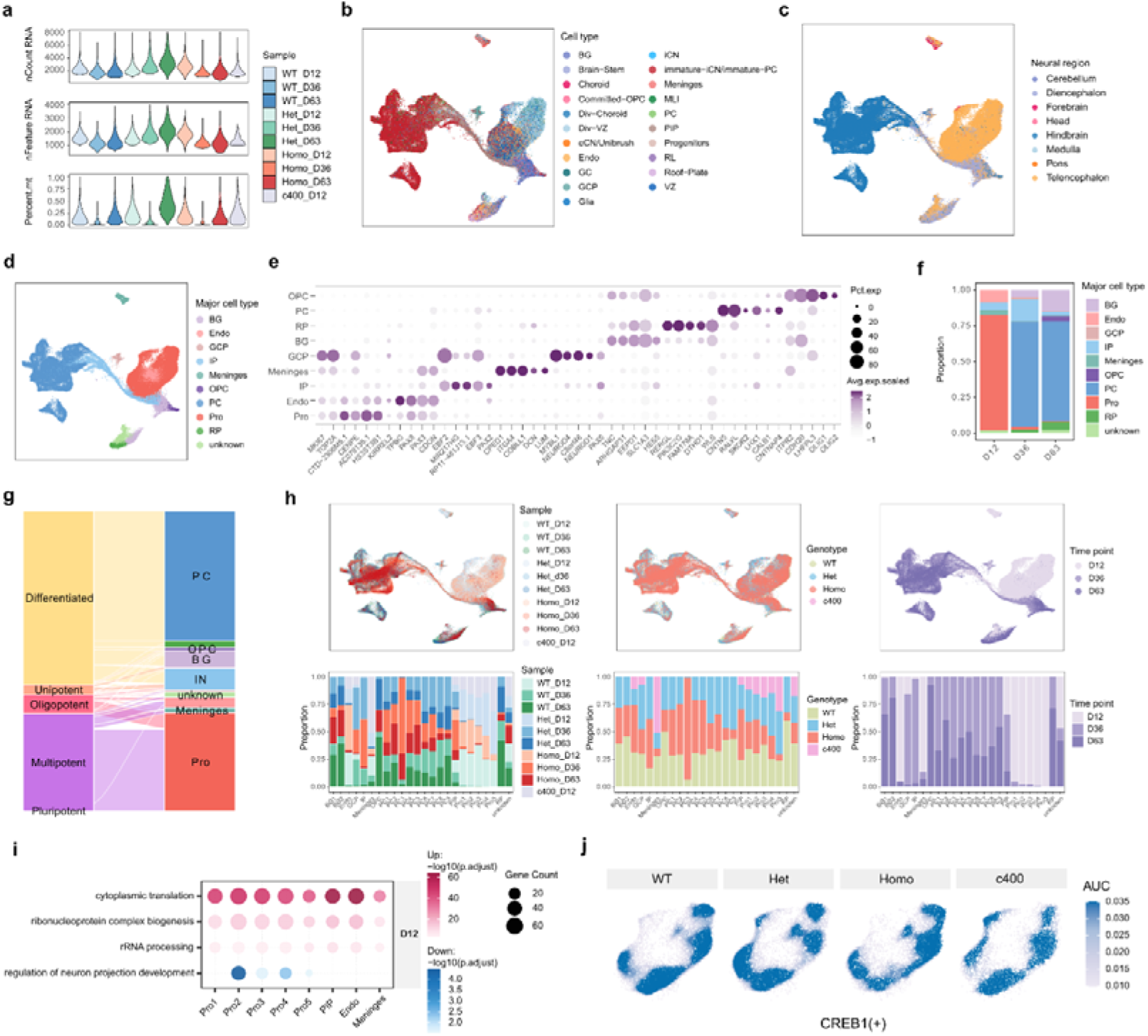
Temporal transcriptomic alterations in TDP2-mutant hCerOs. **a,** Violin plots displaying the number of counts, the number of genes, and the mitochondrial ratio for each sample. **b,** UMAP plot showing the label transfer results from a published human cerebellar organoid scRNA-seq reference dataset, coloured by reference cell type. **c,** UMAP plot showing the label transfer results from a published human neural organoid snRNA-seq reference dataset, coloured by reference neural regions. **d,** UMAP visualization showing the characterized major cell types. **e,** Dot plot displaying the marker-gene expression level across major cell types. **f,** Bar plot comparing the composition of major cell types by time point. **g,** Sankeyplot showing the CytoTRACE developmental-potency categories of all cells. **h,** UMAP visualizations coloured by sample, genotype and developmental time point (top). Bar plot comparing the composition of cellular subtypes by sample, genotype, and developmental time point (bottom). **i,** Dot plots showing GO terms enriched in DEGs in Homo versus Het at D12. **j,** Violin plots showing CREB1(+) regulon activity, quantified by AUCell, in progenitor populations of each genotype at D12.

**Extended Data Fig. 5.**
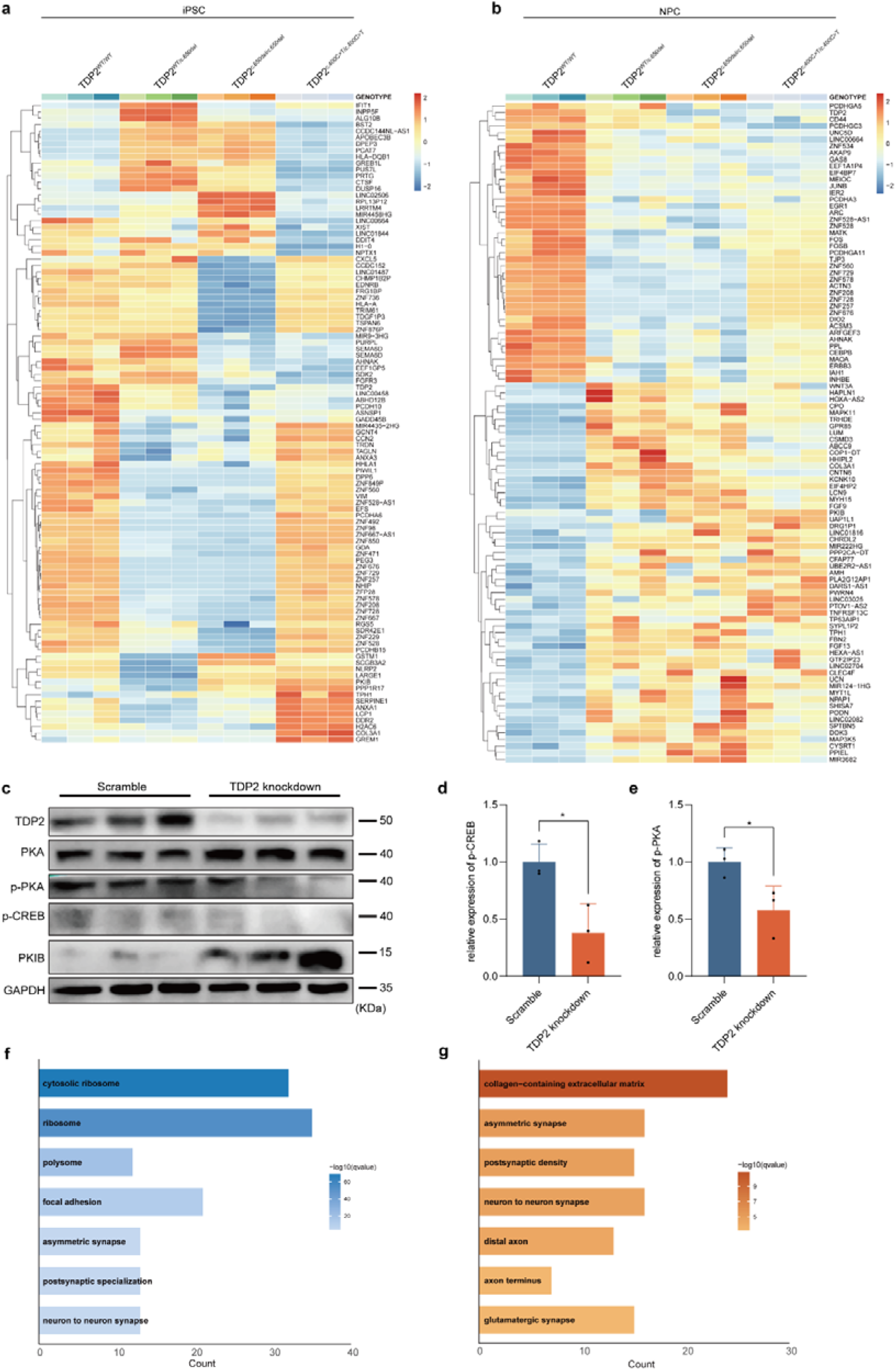
RNAseq analysis of iPSCs and iNPCs. **a,b,** Heatmaps showing gene expression profiles of iPSCs **(a)** and iNPCs **(b)** from bulk RNA-seq data. **c-e,** Representative western blots of TDP2 knockdown iPSCs by shRNA **(c)**. Quantitative analysis of p-CREB/CREB and p-PKA expression **(d,e)**. **f,g,** Barplots showing enriched GO terms for common downregulated **(f)** and upregulated **(g)** genes in TDP2 c.650del and c.400C>T mutant iNPCs versus WT TDP2 iNPCs. (mean ± SD, n = 3). Statistical test: one-way ANOVA. *\*\*\*\*P* < 0.0001, *\*\*\*P* < 0.001, *\*\*P* < 0.01, *\*P* < 0.05.

**Extended Data Fig. 6.**
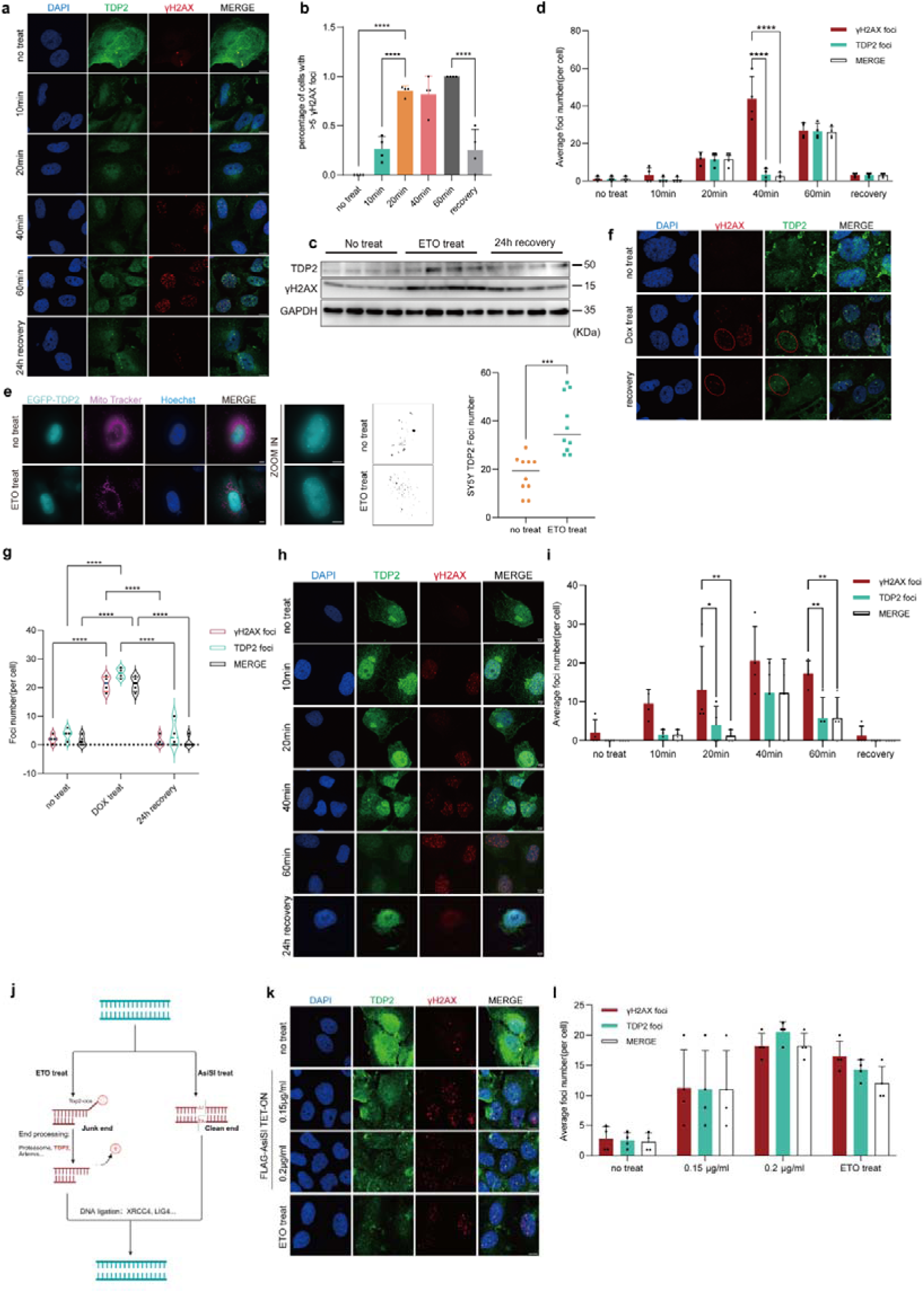
Formation and proximity of TDP2 and γH2AX foci in response to etoposide-induced DNA damage. **a,** Representative immunofluorescence of DNA damage responses in SH-SY5Y cells treated with etoposide (ETO) across a 60-minute induction and 24-hour recovery phase. Scale bars, 10 μm. **b,** Quantification of the percentage of γH2AX-positive SH-SY5Y cells after ETO treat (Mean intensity ± SD n = 4). **c,** Representative of western blot for TDP2 expression during acute ETO-induced DNA damage and subsequent recovery. **d,** Quantification of TDP2 foci and distance to γH2AX foci showing MERGE (foci are less than 5 pixels, being 0.36 μm, apart). Data are from n = 4 biological replicates counting about 40 cells for every time point. **e,** Live-cell super-resolution microscopy of SH-SY5Y cells expressing EGFP-TDP2. Scale bars, 5 μm. Scale bars in zoom in figure, 2.5 μm. Quantification of TDP2 foci following ETO treatment (n = 10, counting about 30 cells for control and ETO-treated respectively). Scale bars, 5 μm. Scale bars in zoom in figure, 2.5 μm. **f,** Representative immunofluorescence images of SH-SY5Y cells after doxorubicin (DOX) treatment and 24h recovery. Scale bars 15.1 μm. **g,** Quantification of the number of TDP2 foci and γH2AX foci overlapping in SH-SY5Y cells after DOX treatment and recovery. Data are from n = 4 biological replicates, counting about 40 cells for each treatment. **h,** Representative immunofluorescence images of SH-SY5Y cells after 60min camptothecin (CPT) treatment and 24h recovery. Scale bars, 5 μm. **i,** Quantification of the number of TDP2 foci and γH2AX foci overlapping in SH-SY5Y cells after CPT treatment and recovery. Data are from n = 4 biological replicates, counting about 40 cells for each treatment. **j,** Schematic of the double-strand breaks (DSBs) repair process for “clean end” damage and “junk end” damage in cells. **k,l,** SH-SY5Y Cells transfected with FLAG-AsiSI and treated (0.15μg/ml, 0.2μg/ml) or not treated with tetracycline, which induces the ‘clean’ genome-wide sequence-specific DSBs. Scale bars, 10 μm **(k)**. Quantification of TDP2 foci and distance to γH2AX foci showing overlapping and distant. Data are from n = 4 biological replicates counting about 40 cells for every treatment **(l)**. Statistical test: one-way ANOVA or two-way ANOVA. *\*\*\*\*P* < 0.0001, *\*\*\*P* < 0.001, *\*\*P* < 0.01, *\*P* < 0.05.

**Extended Data Fig. 7.**
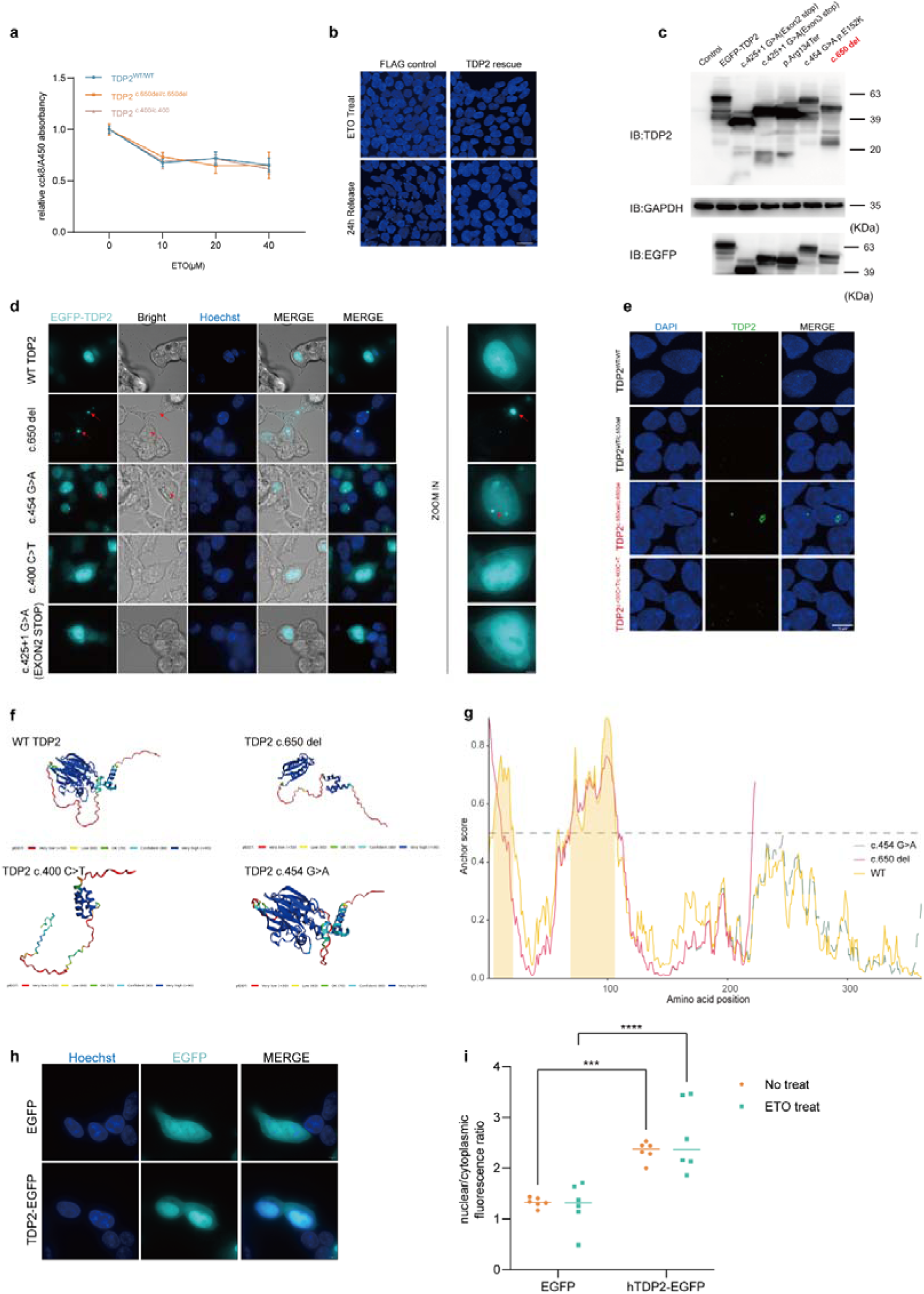
SCAR23-associated TDP2 mutations drive aberrant nuclear condensation. **a,** Quantification of cell viability after ETO treat in TDP2 WT and mutant iPSCs (mean ± SD, n = 4). b, Representative Immunofluorescence images of FLAG and FLAG-TDP2 overexpression in TDP2 c.650del iPSC cells after ETO treatment and 24h recovery. Scale bars, 25 μm. c,d, Representative of live cell immunofluorescence and Western blot images for exogenous expression of different SCAR23 TDP2 mutant genes with EGFP tag in 293T cells. Scale bars, 5 μm, zoom in scale bars 2 μm. e, Representative Immunofluorescence images of pre-permeabilized iPSCs. f,g, structures of WT and mutant TDP2 predicted by AlphaFold 3. Colours ranging from blue to red depict the per-residue measure of local confidence (pLDDT) for the model (f). Disorder analysis of WT and mutant TDP2 sequences using AlphaFold 3 Metapredict scores (g). h, Representative of live cell immunofluorescence for exogenous expression of d C-terminally tagged TDP2-EGFP in 293T cells. Scale bars, 5 μm. i, Quantification of the TDP2-EGFP intensity ratios of nuclear/cytoplasmic after ETO treatment normalized to EGFP control (Mean intensity ± SD, n = 6). Statistical test: two-way ANOVA. *\*\*\*\*P* < 0.0001, *\*\*\*P* < 0.001.

**Extended Data Fig. 8.**
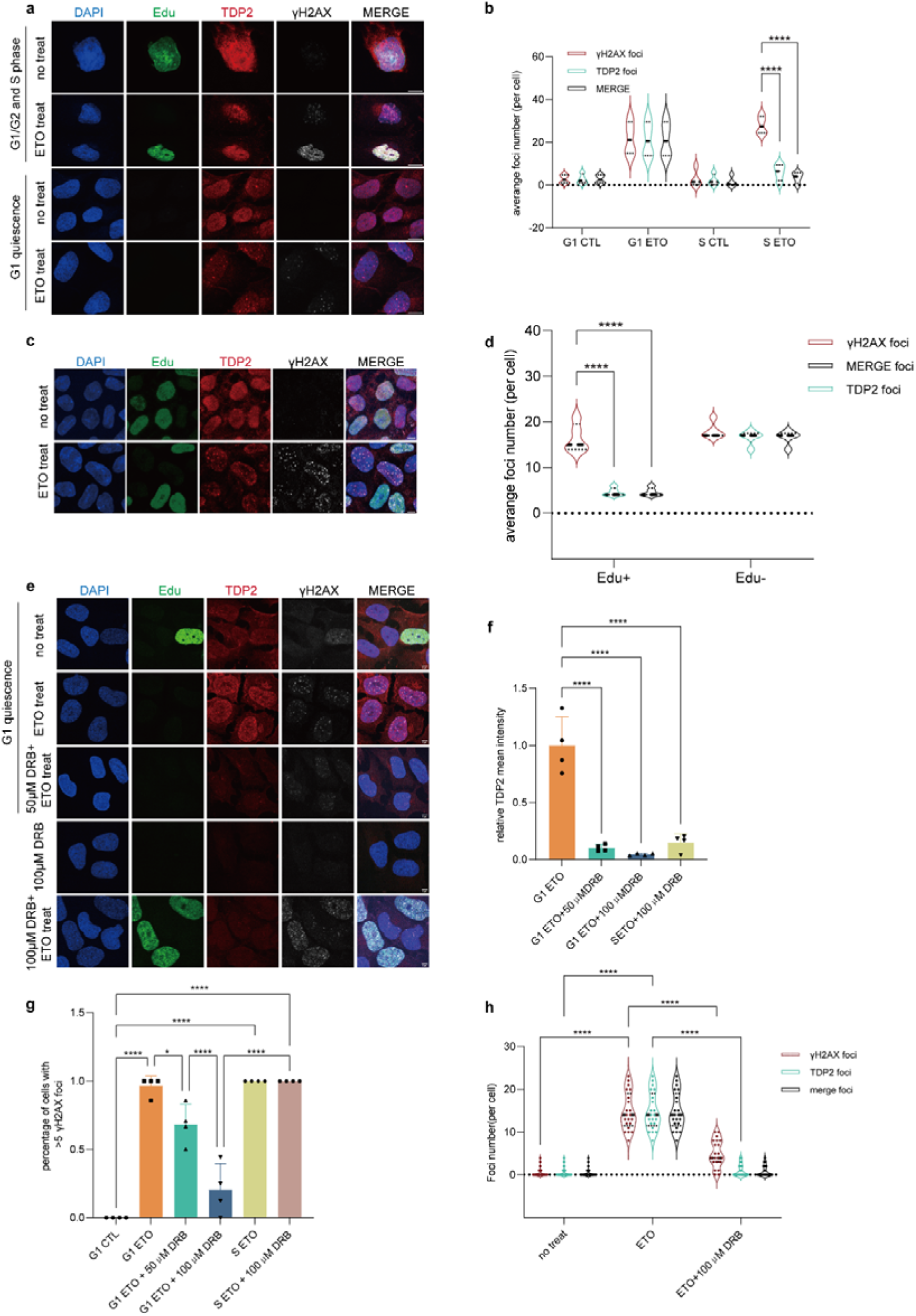
TDP2 Foci condense at transcription-coupled DSBs site in G1 phase. **a,** Representative immunofluorescence images of SH-SY5Y cells after G1 quiescence and ETO treat. Scale bars, 10 μm. **b,** Quantification analysis of TDP2 and γH2AX foci in SH-SY5Y cells in different cell phase after ETO treat. Data are from n = 4 biological replicates, counting about 40 cells for each treatment. **c,** Representative immunofluorescence images of human NPC cells with ETO treat. Scale bars 10 μm. **d,** Quantification analysis of TDP2 and γH2AX foci in NPC cells with ETO treatment. Data are from n = 5 biological replicates, counting about 50 cells for each treatment. **e,** Representative immunofluorescence images of SH-SY5Y cells in G1 quiescence and ETO treatment with transcription inhibition by DRB. Scale bars 10 μm. **f,g,** Quantification analysis of TDP2 intensity **(f)** and percentage of γH2AX positive cells **(g)** of SH-SY5Y cells after DRB and ETO treatment in G1 and G2/S phase (Mean intensity ± SD, n = 4). **h,** Quantification analysis of TDP2 and γH2AX foci in SH-SY5Y cells with DRB and ETO treatment. Data are from counting about 21 cells in 3 biological replicates for each treatment. Statistical test: one-way ANOVA (f,g) or two-way ANOVA (b,d,h). *\*\*\*\*P* < 0.0001, *\*\*\*P* < 0.001, *\*\*P* < 0.01, *\*P* < 0.05.

**Extended Data Fig. 9.**
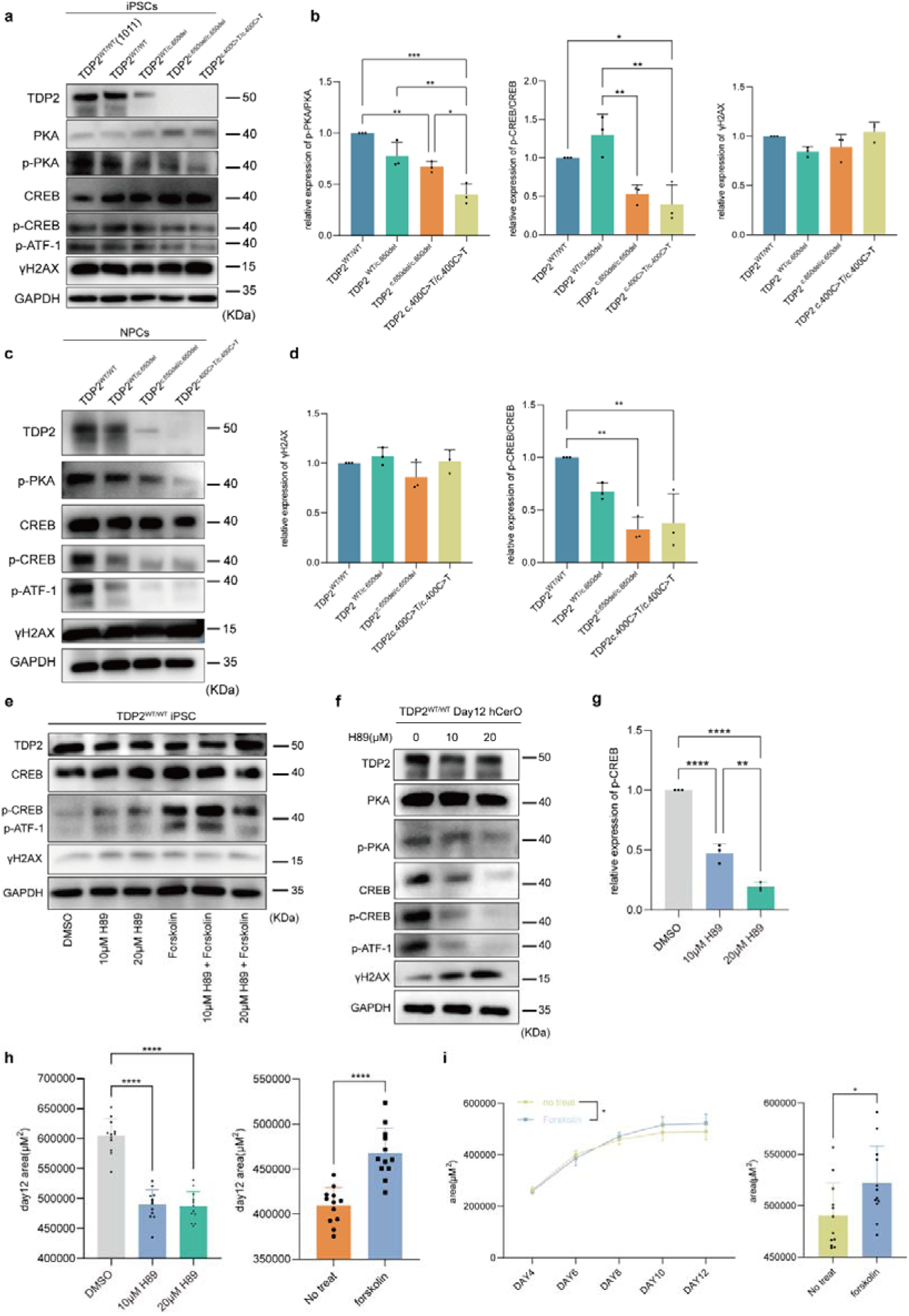
Inhibition and activation of PKA–CREB axis in iPSCs, iNPCs and hCerOs. **a,b,** Representative western blot of iPSCs for PKA–CREB pathway protein and γH2AX. Quantification of band intensities for p-PKA/PKA, p-CREB/CREB, γH2AX in iPSCs, ratios normalized to control (mean ± SD, n = 3). **c,d,** Representative western blot analysis of iNPCs for PKA–CREB pathway protein and γH2AX **(c)**. Quantification of band intensities for p-CREB/CREB and γH2AX, ratios normalized to control **(d)** (mean ± SD, n = 3). **e,** Representative western blot of iPSCs with H89 or forskolin treatment. **f,g,** Representative western blot of day 12 hCerOs with H89 treatment **(f)**. Quantification of band intensities for p-CREB/CREB ratios normalized to DMSO control **(g)** (mean ± SD, n = 3). **h,** Quantitative analysis of day 12 hCerOs area of WT TDP2 line with H89 treatment and TDP2 c.650del line with forskolin treatment (mean ± SD, n = 12). **i,** Quantitative analysis of hCerOs area of TDP2 c.400C>T line with forskolin treatment from day4 to day 12 (mean ± SD, n = 12). Statistical test: one-way ANOVA (b,d,g,h) and two-way ANOVA (i). *\*\*\*\*P* < 0.0001, *\*\*\*P* < 0.001, *\*\*P* < 0.01, *\*P* < 0.05.

**Extended Data Fig. 10.**
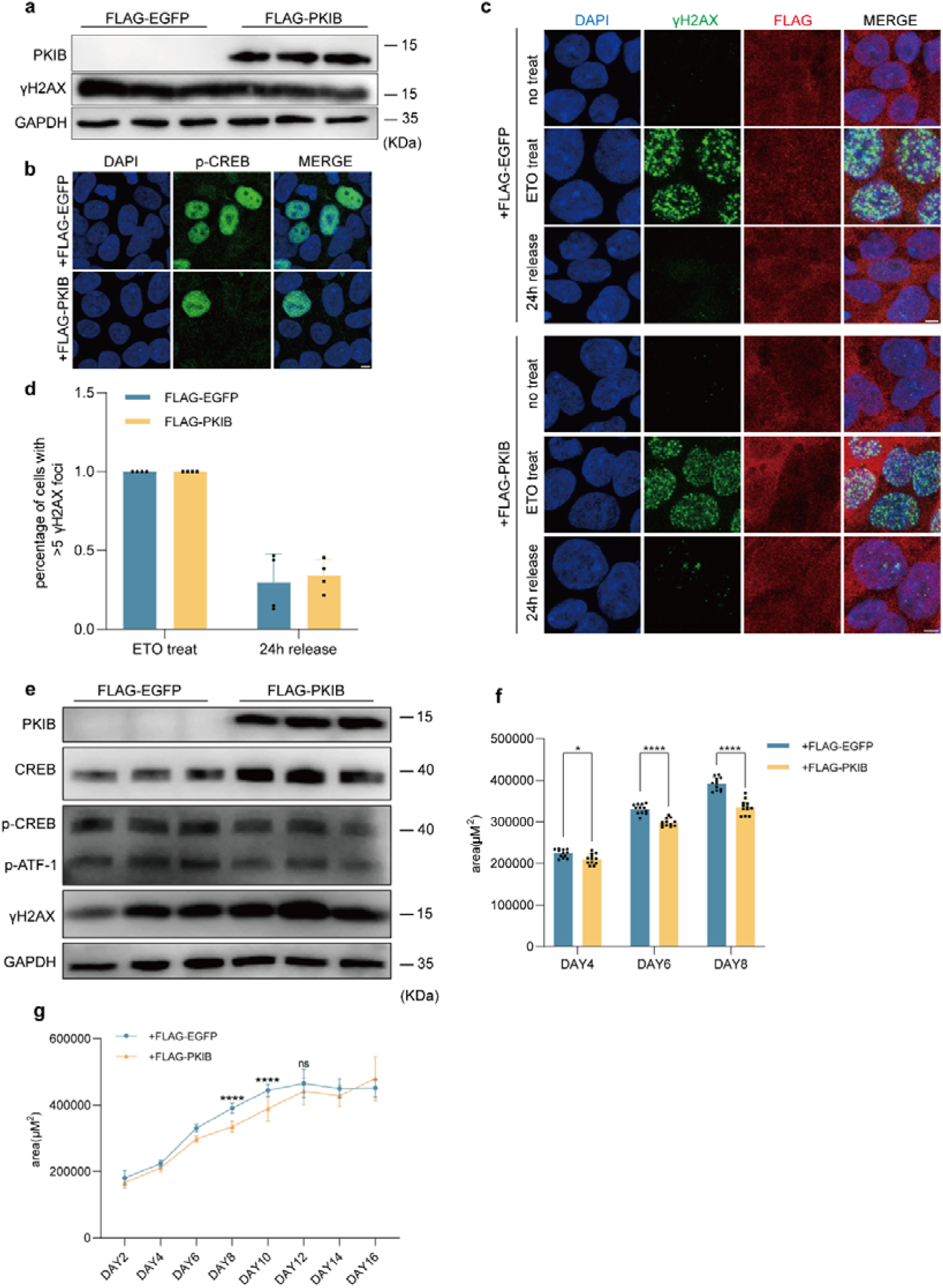
PKIB overexpression cause DNA damage accumulation and hypoplasia during hCerOs development. **a,** Representative western blot of iPSCs for PKIB and γH2AX with FLAG-EGFP or FLAG-PKIB overexpression. **b,** Representative immunofluorescence images of p-CREB for iPSCs. Scale bars 5 μm. **c,d,** Representative immunofluorescence images and quantification of γH2AX and anti-FLAG antibody for iPSCs. Scale bars 5 μm. (mean ± SD, n = 4). **e,** Representative western blot of day 12 hCerOs with FLAG-EGFP or FLAG-PKIB overexpression. **f,g,** Quantitative analysis of area of hCerOs with FLAG-EGFP and FLAG-PKIB overexpression from day 2 to day 12 (mean ± SD, n = 12). Statistical test: two-way ANOVA. *\*\*\*\*P* < 0.0001, *\*\*\*P* < 0.001, *\*\*P* < 0.01, *\*P* < 0.05.

**Extended Data Fig. 11.**
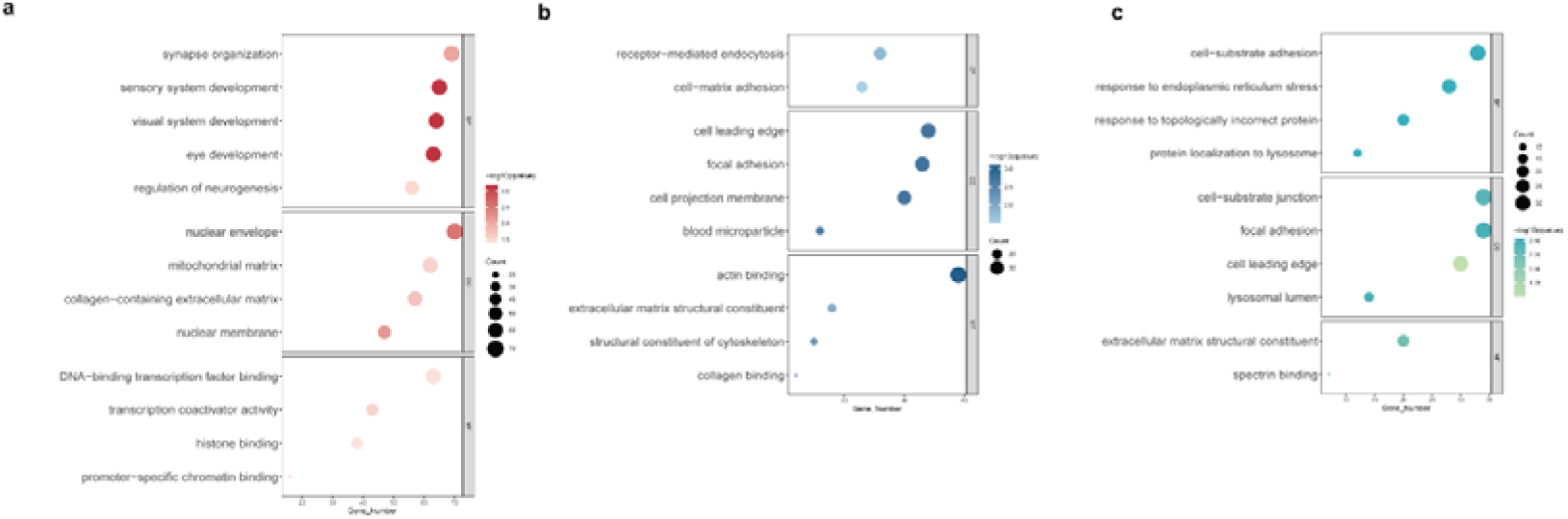
GO enrichment analysis of SCAR23 WGS. **a-c**, GO enrichment analysis of unique 1-4bp deletion mutation in the homozygous TDP2 c.650del mutation (Homo) **(a)**, healthy sibling (WT) **(b)**, heterozygous TDP2 c.650del mutation genome **(c)**.

**Extended Data Fig. 12.**
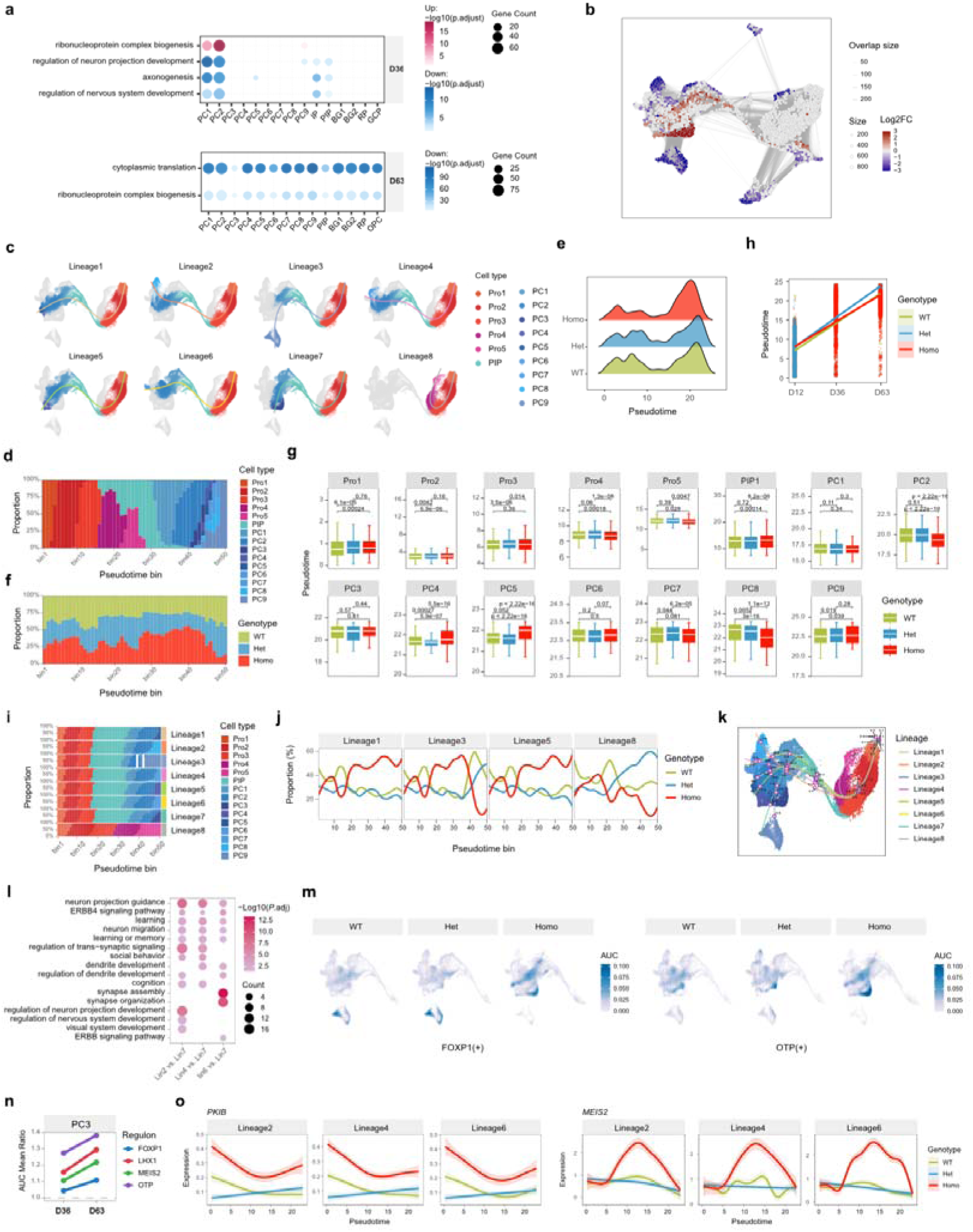
TDP2 inactivation alters the developmental trajectory of major neuronal lineages and perturbs their regulatory networks. **a,** Dot plots showing broadly enriched GO terms of DEGs in Homo versus Het hCerOs at D36 (top) and D63 (bottom). **b,** Milo neighbourhood graph visualizing differential abundance between disease (Homo cells) and control (WT and Het cells). **c,** Visualization of eight individual Slingshot reconstructed lineages, cells coloured by cellular subtype, curves coloured by lineage. **d,f,** Cellular subtype **(d)** and genotype **(f)** proportions along 50 pseudotime bins. **e,** Density plot showing pseudotime distribution in WT, Het, and Homo cells. **g,** Boxplot showing cell type-specific pseudotime distributions for representative neuronal cells across genotypes. **h,** Relationship between chronological culture time (D12, D36, and D63) and pseudotime for each genotype. **i,** Composition of cell type across pseudotime bins in each lineage, highlighting the late-pseudotime representation of PC3 in Lineage 7. **j,** Genotype proportions along pseudotime bins in four selected Slingshot lineages (Lineages 1, 3, 5, and 8). **k,** Slingshot trajectories with fitted knots labelled. **l,** Bubble plot showing enriched GO terms for genes distinguishing the Homo-enriched Lineage 7 from WT-enriched Lineages 2, 4, and 6 within the knot 4-5 interval. **m,** UMAP visualization of FOXP1(+) and OTP(+) regulons activity across genotypes. **n,** Homo-to-WT ratios of mean regulon activity scores for MEIS2(+), LHX1(+), FOXP1(+), and OTP(+) in PC3 at D36 and D63. **o,** Smoothed expression trajectories of *PKIB* and *MEIS2* in Lineages 2, 4 and 6, stratified by genotype.

**Extended Data Fig. 13.**
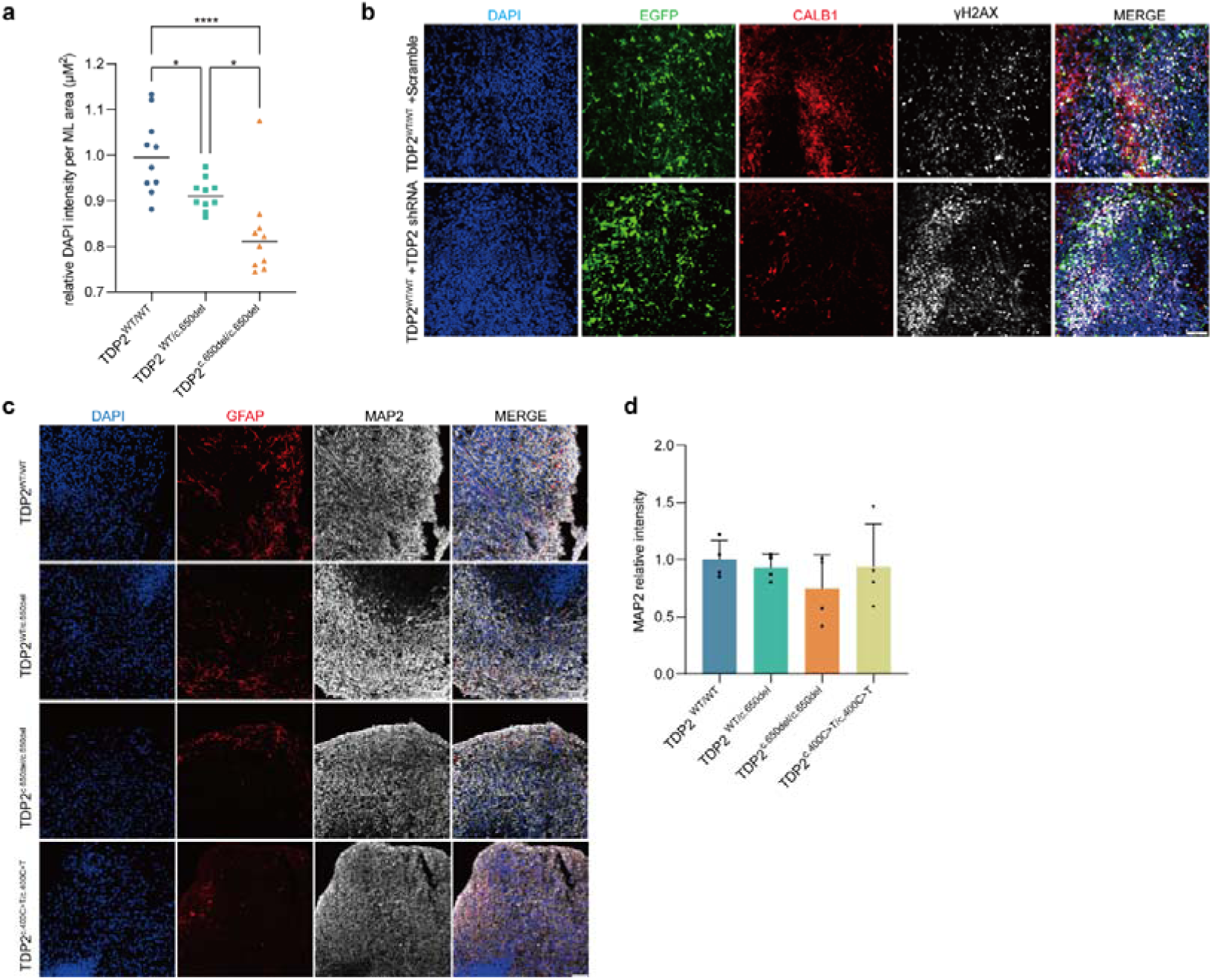
MAP2 and GFAP expression pattern in TDP2 mutant and CALB1 expression in TDP2 knockdown hCerOs. **a,** Quantification of DAPI intensity per area in ML/PL of day 60 hCerOs (mean intensity ± SD, n = 10). **b,** Representative immunofluorescence images of CALB1 and γH2AX for day 60 TDP2 knockdown hCerOs by shRNA. Scale bars, 50 μm. **c,** Representative immunofluorescence images of GFAP and MAP2 for day 100 TDP2 mutant hCerOs. Scale bars, 100 μm. **d,** Quantification analysis of MAP2 relative intensity for day 100 hCerOs with TDP2 mutation (mean intensity ± SD, n = 4).

**Extended Data Fig. 14.**
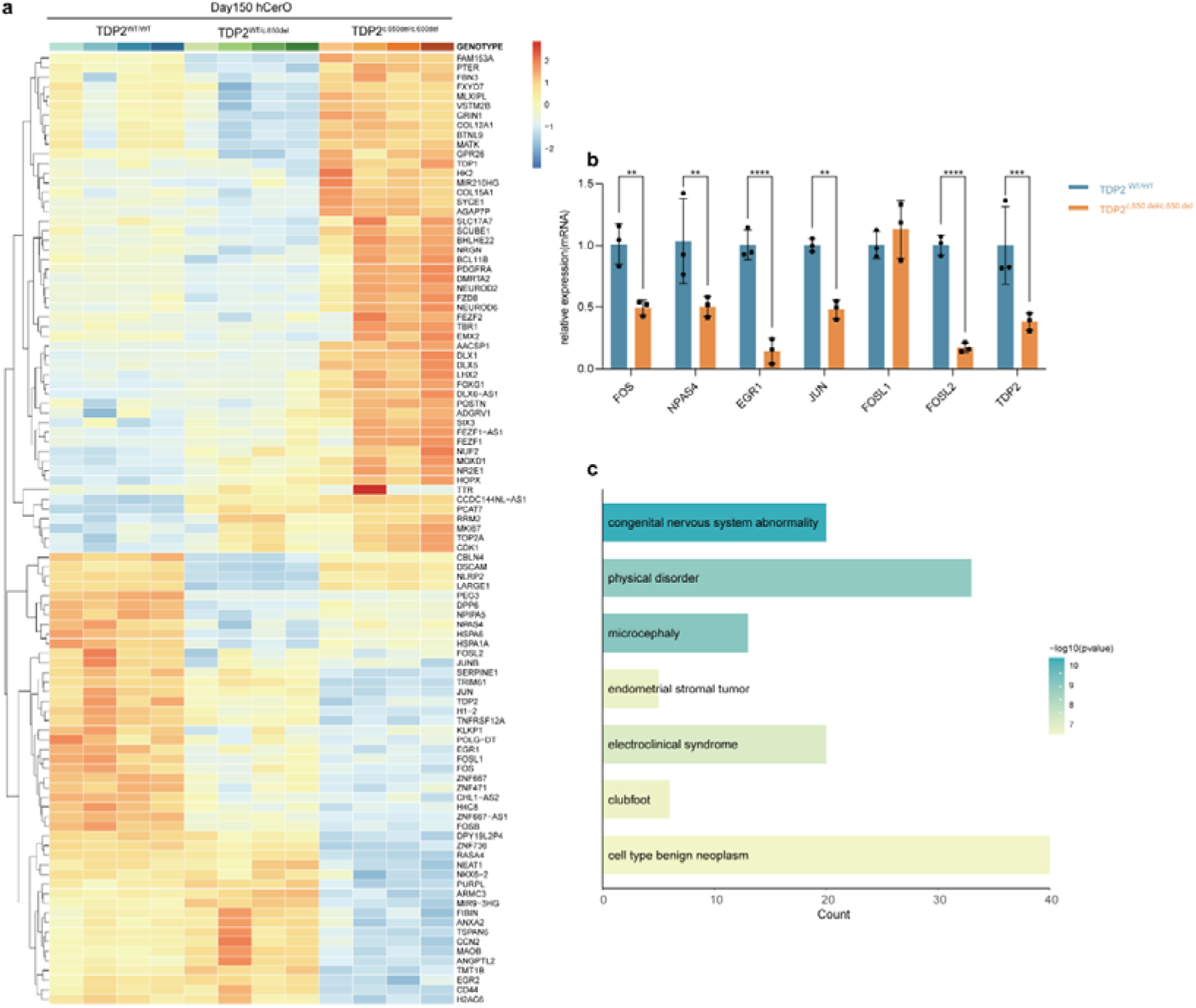
ERGs down regulation in TDP2 mutant day 150 hCerOs. **a,** Heatmap analysis of bulk RNA-seq data for day 150 hCerOs. **b,** Quantification of the expression of TDP2 and ERGs in day 150 hCerOs using RT-qPCR (mean ± SD, n = 3). **c,** Disease enrichment analysis of differential expression genes in TDP2 c.650del mutant hCerOs compared to WT. Statistical test: two-way ANOVA. *\*\*\*\*P* < 0.0001, *\*\*\*P* < 0.001, *\*\*P* < 0.01, *\*P* < 0.05.

**Extended Data Fig. 15.**
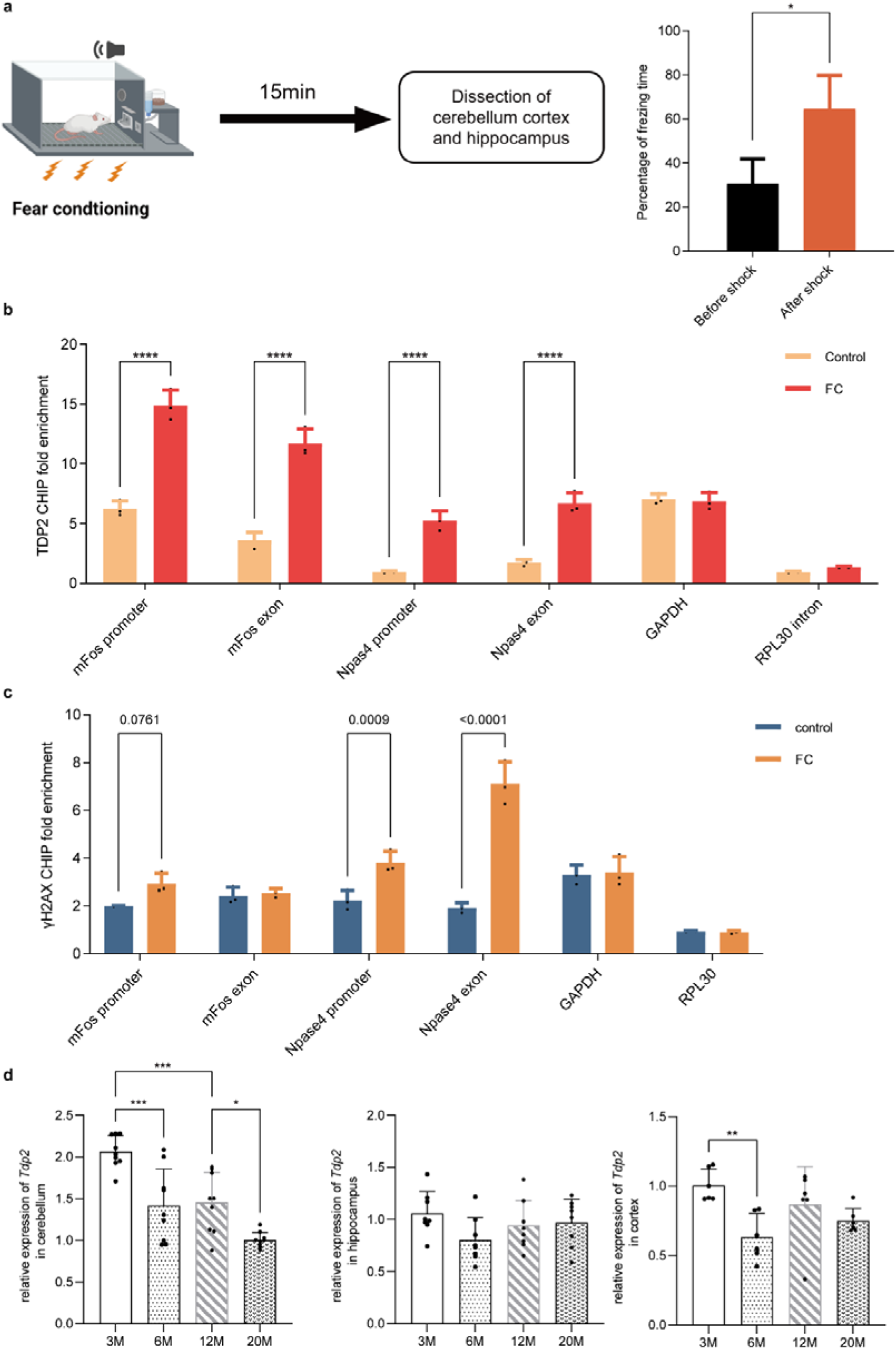
Tdp2 expression and binding in mouse after fear conditioning. **a,** Two-month-old C57BL/6 mice were placed in a contextual chamber for 2 min, following which two 1 mA foot shocks were delivered from the grid floor. The mice remained in the chamber for an additional 30s and were then moved back to their home cages. Fifteen minutes following the footshock, animals were measured for freezing time in chamber following which the cortex, hippocampus, and cerebellum were sacrificed for CHIP-qPCR (mean ± SD, n = 3). **b,c,** ChIP qPCR analysis of Tdp2 **(b)** and γH2AX **(c)** binding to the promoters of *Fos* and *Npas4* in the mouse cerebellum after fear conditioning. RPL30 intron and promoters of GAPDH were probed as two negative controls (mean ± SD, n = 3). **d,** Quantitative analysis of *Tdp2* expression in cerebellum, hippocampus and cortex mice from age of 3 months (3M) to 20 months (20M) by qPCR (mean ± SD, for cerebellum and hippocampus n = 8, for cortex n = 6). Statistical test: one-way ANOVA (d) and two-way ANOVA (b,c). *\*\*\*\*P* < 0.0001, *\*\*\*P* < 0.001, *\*\*P* < 0.01, *\*P* < 0.05.

**Extended Data Fig. 16.**
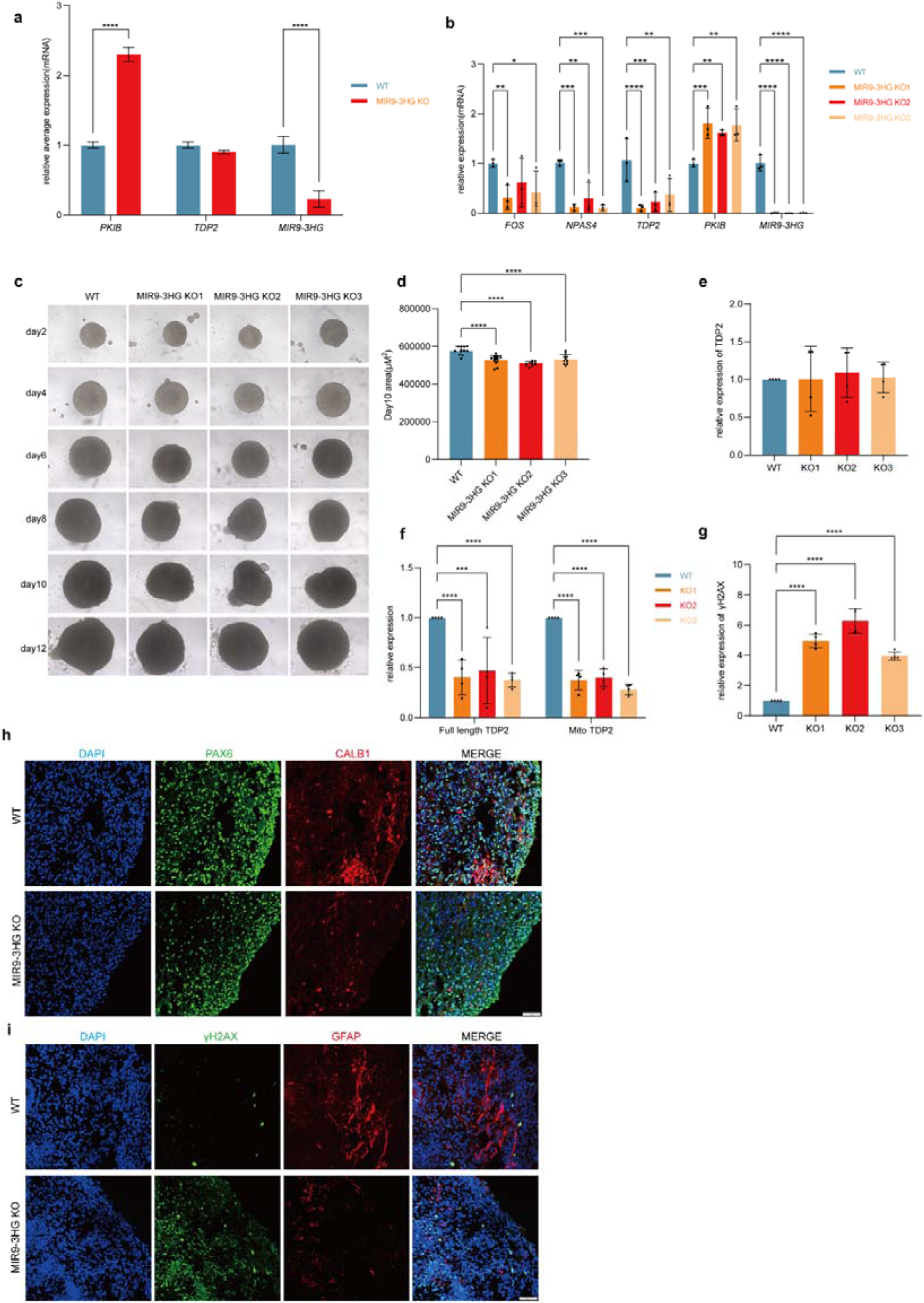
MIR9-3HG knockout cause DNA damage repair deficiency in cerebellar development. **a,** Quantification of the *TDP2, PKIB* and *MIR9-3HG* expression in MIR9-3HG KO iPSCs using RT-qPCR (mean ± SD, n = 3). **b,** Quantification of the *TDP2* and *ERGs* expression in MIR9-3HG KO day 150 hCerOs using RT-qPCR (mean ± SD, n = 3). **c,d,** Representative brightfield images and quantitative analysis area of MIR9-3HG KO hCerOs (d, Scale bars, 200 μm). Quantitative analysis of hCerOs area from day 2 to day 12 (e,f) (mean ± SD, n = 12). **e-g,** Quantification of the expression of full TDP2 and mitochondria TDP2 for iPSCs **(e)** and day 100 hCerOs **(f)** of MIR9-3HG KO lines ratios normalized to WT control (n = 4). Quantification of γH2AX level in MIR9-3HG knockout day 150 hCerOs ratios normalized to WT control (n = 4) **(g)**. **h,i,** Representative immunofluorescence images of PAX6 / CALB1 and γH2AX / MAP2 intensity for day 60 WT and MIR9-3HG KO hCerOs (mean ± SD, n = 4). Scale bars 50 μm. Statistical test: one-way ANOVA (d,e,g) and two-way ANOVA (a,b,f). *\*\*\*\*P* < 0.0001, *\*\*\*P* < 0.001, *\*\*P* < 0.01, *\*P* < 0.05.

